# CFD-Informed Hybrid Modeling Unlocks Scalable, Tunable Amino Acid Production in *Methanothermobacter marburgensis*

**DOI:** 10.64898/2026.07.09.737395

**Authors:** Benedikt Haslinger, Barbara Reischl, Franziska Steger, Maximilian Krippl, Lukas Gsenger, Elisa Hilts, Aquilla Ruddyard, Michael Stadlbauer, Stefan Drießler, Hayk Palabikyan, Günther Bochmann, Mark Duerkop, Simon K.-M. R. Rittmann

## Abstract

Methanogenic archaea, such as Methanothermobacter marburgensis, represent a powerful biological platform for carbon capture and valorization, directly converting carbon dioxide (CO2) and molecular hydrogen (H2) into proteinogenic amino acids (AAs). In this study, we present a controlled and scalable strategy for tailoring AA production (biosynthesis and secretion) in continuous gas fermentation. By applying various Design of Experiments (DOE) techniques, we systematically identified and optimized key process parameters governing AA biosynthesis and shaping a targeted AA secretion profile. A hybrid modeling framework combining experimental data with scale-independent parameters derived from computational fluid dynamics (CFD) enabled robust performance prediction across bioreactor scales. This model-driven approach successfully translated the process from 120 mL glass bottles via 2 L to 150 L reactors, corresponding to a reaction-volume scale-up factor of 2000. These findings set the foundation for a robust and predictive platform for sustainable AA production, positioning archaea as a high-potential alternative in industrial biotechnology.

## 1. Introduction

The worldwide market for proteinogenic amino acids (AAs) is growing steadily and the demand is expected to increase annually by about 8.3% from 2024–2034^1,2^, most notably in the animal feed sector^3^ as well as functional food ingredients or pharmaceuticals^4^. The AA market has become a branch of biotechnology that is now worth several billion USD^1,3,5^. Fermentation has been used for AA production for more than 50 years^6^. The most common bacterial AA overproducing organisms are *Escherichia coli* and *Corynebacterium glutamicum*, which are among the established microbial cell factories for industrial production of single AAs^5,7,8^. To reach a desired yield or AA profile, the current system of choice is to genetically engineer an organism and readjust its metabolic pathways^9^. As the focus and demand for specialized AA mixtures continues to grow, new research is needed to provide the increasing market needs with tailored biotechnological solutions^3,10^.

Archaea are increasingly important to biotechnological applications due to their ability to withstand harsh and extreme environments and their unique metabolic pathways, which can be used for the biofuel production, high-value chemicals, and AAs^11–13^. Methanogenic archaea (methanogens) are anaerobic microorganisms that are best known for their ability to generate methane (CH_4_) by reducing carbon dioxide (CO_2_) with molecular hydrogen (H_2_)^14^ – a process that has been extensively studied for its potential in renewable CH_4_ production^15–22^. Over the past years, methanogens have become model organisms for a variety of studies for astrobiology^23–25^, geobiology^26^, ecophysiology^24,27^ and biotechnology^18,21,28–31^. Recently, methanogens have been shown to secrete up to 20 AAs into the supernatant simultaneously^32,33^. The ability of methanogens to produce a broad spectrum of AAs from CO_2_ and H_2_ warrants further exploration to utilize them as a potential platform for sustainable AA production, as the current bacterial AA overproducers, *E. coli* and *C. glutamicum*, are specialized in producing only single AAs and use sugar as feedstock^3,5,7,34^. Therefore, using H_2_ and CO_2_ as substrates and the methanogen *Methanothermobacter marburgensis* for AA production^35–39^ could offer an advantage for developing and commercializing a scalable and sustainable AA production bioprocess.

Realizing this potential, however, requires systematic process development to identify and optimize key drivers of performance. To investigate and optimize bioprocesses towards a certain output, regarding critical process parameters (CPPs) as well as media compositions, design of experiments (DOE) has been widely employed for various microorganisms since the late 20^th^ century^40,41^. More recently, intensified DOE (iDOE) approaches have emerged as powerful tools that enable more efficient gathering of data for understanding process dynamics and the impact of parameters on a bioprocess. In contrast to traditional DOE – where each experiment is typically performed under static conditions – iDOE introduces deliberate, time-dependent changes of one or more CPPs within the same experimental run^42,43^. For instance, instead of maintaining a constant feed rate or temperature throughout a cultivation, iDOE involves stepwise changes in these parameters within a single run, enabling observation of the system’s dynamic response to a broader range of conditions. These intra-experimental shifts allow for extensive exploration of the process parameter space within fewer experiments, significantly reducing time and resource requirements while generating rich, time-resolved data. Both DOE and iDOE offer methodologies for experimental design and data generation, however meaningful insights can only be created using appropriate modeling techniques. Classical DOE approaches, like the response surface methodology (RSM), are still widely used for endpoint-data, e.g., for biomethanation^21^, but fall short in interpreting data generated using dynamic shifts in CPPs or describing bioprocesses in a time-resolved manner^41^. Regardless of the employed experimental design method, the combination of a data-driven framework with mechanistic constraints and process understanding to so-called hybrid models has proven to facilitate more accurate predictions while lowering the required experimental effort across various up-and downstream unit operations^42,44–47^. Beyond process optimization and characterization, the scalability of developed processes and their respective models remains a key challenge in bioprocess development. Utilizing scale-independent parameters derived from computational fluid dynamics (CFD) is a key enabler for the effective implementation of hybrid models in process development. “Scale-independent” refers to CFD-derived descriptors that capture underlying hydrodynamic conditions and can be intentionally matched across scales by appropriate adjustment of scale-specific operating parameters, rather than to operating parameters themselves such as stirrer speed or gas flow rate. Without CFD, optimizations made at laboratory scale often fail to translate successfully to pilot or industrial-scale production due to unaccounted-for changes in the reactor environment such as shear and gassing conditions. CFD provides a framework for understanding and predicting these scale-dependent phenomena, enabling the accurate scale-wise extrapolation of optimized conditions. This capability is critical for ensuring that performance and product quality are preserved throughout scale-up, rendering CFD models a key enabler of successful model across the entire process life cycle^48–52^.

The first objective of this work was to establish two complementary scaling criteria for AA production. The qualitative criterion aimed to tailor the AA profile by adjusting the media composition, in particular trace element (TE) availability, to reduce alanine and glutamic acid – two dominant components of the product spectrum – in order to shift production toward a higher proportion of less abundant AAs (residual AAs). This is relevant for microbial protein applications, where product quality is determined not only by total AA concentration but also by the composition of the AA profile. Additionally, TE availability has been shown to influence enzymatic activity and metabolic routing in methanogenic microorganisms^15,53^. The quantitative criterion was defined as the biomass specific AA yield (Y_AA/x_). These were derived through a novel and unique combination of data-driven and mechanistic modeling as well as CFD simulations. To support scale translation, two scale-independent CFD-derived parameters, the volumetric mass transfer coefficient of H_2_ (k_L_aH_2_) and the average shear rate (*γ̇_avg_*) in the reactor, were selected as additional hybrid model inputs, with *γ̇_avg_* serving as a hydrodynamic descriptor to complement k_L_aH_2_ in capturing scale-dependent mixing and gas dispersion effects. The second objective was to transfer both criteria across scales, from 120 mL closed batch in serum bottles to continuous cultures in 2 L and 150 L bioreactors. To achieve these objectives, the study strategically combined various DOE, iDOE and modeling approaches to enable a time-efficient and robust transfer of bioprocess performance and AA profiles across scales.

## 2. Results

### 2.1. Media optimization

To reduce the formation of unwanted AAs during continuous production, a series of closed batch experiments were performed using the described TE screening design. Therefore, TE concentrations were systematically varied to evaluate whether they influence the AA profile. The DOE results demonstrated that specific TE combinations affected the relative AA composition and enabled the optimization that shifted the spectrum toward the desired profile. This shift was targeted to reduce the dominance of alanine and glutamic acid and thereby increase the relative contribution of other AAs within the spectrum. The resulting data was used to train a predictive model and optimize for reduced alanine and glutamic acid. Guided by the model’s predictions, a modified TE formulation was designed with reduced concentrations of cobalt, molybdenum and magnesium, while increasing nickel and maintaining the level of iron (Supplementary table 1). To validate this optimized composition, a continuous cultivation at 2 L scale was conducted and compared to the performance of a reference run using the original TE composition, under otherwise identical process conditions (Figure 1). Following an initial adaption phase during the transition from fed-batch to continuous operation, the optimized TE solution led to a substantial decrease in alanine and glutamic acid production. During the stable production phase alanine was reduced from 29.2% (±1.9% SD) to 16.8% (±0.8% SD) on average (p < 0.001), while glutamic acid production was decreased by 44.7%, from 45.8% (±5.0% SD) to 1.1% (±1.1% SD) (p < 0.001), which resulted in a shift of residual AA production from 25.0% (±6.2% SD) to 82.0% (±1.7% SD) (p < 0.001). The complete AA compositions before and after TE optimization are provided in Supplementary figure 1. While the optimization primarily targeted AA composition rather than total titer, the modified TE formulation resulted in a moderate decrease in total AA concentration from 283 ± 47 mg L^−^¹ to 211 ± 14 mg L^−^¹. Importantly, this reduction was accompanied by a markedly lower variability during steady-state operation, indicating a more stable production regime. Statistical analysis confirmed that these compositional differences significantly exceeded the within-run process variability observed during steady-state operation. The optimized TE solution was then used in all iDOE2 runs as well as the scale-up experiment.

**Figure 1:**
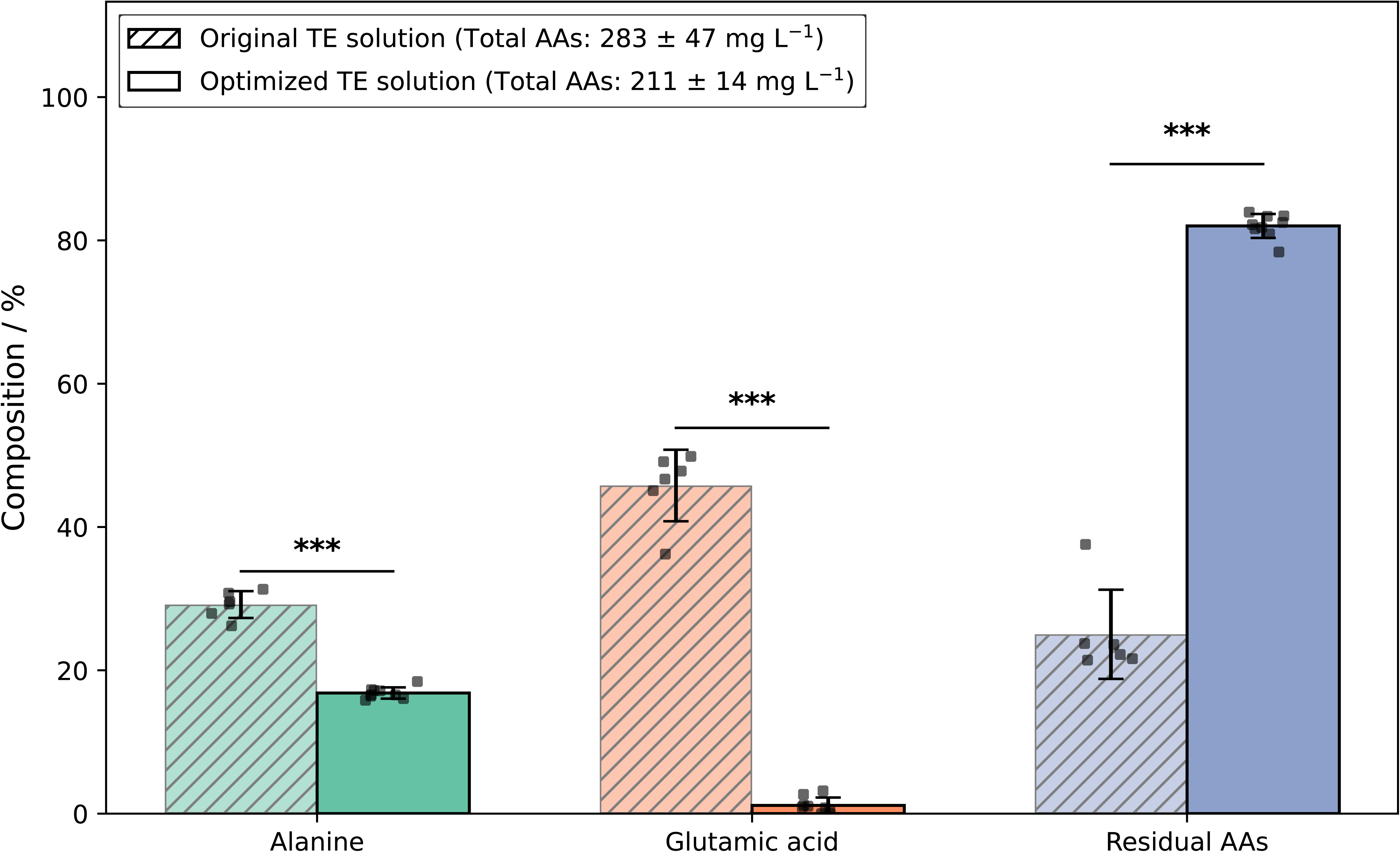
Comparison of AA composition between the original and optimized TE solution during steady-state continuous production. Bars indicate the mean relative composition of alanine, glutamic acid, and the combined fraction of residual AAs, calculated from time-resolved samples collected during the steady-state phase. Individual points represent all time-resolved steady-state measurements. Error bars denote the standard deviation of these measurements within each condition. Statistical significance between original and optimized TE solutions was assessed using two-sided Student’s t-tests on time-resolved steady-state samples (*** p < 0.001). The optimized TE solution resulted in pronounced reductions in alanine and glutamic acid, accompanied by a corresponding increase in the relative proportion of residual amino acids. The complete AA compositions are provided in Supplementary figure 1.

### 2.2. iDOE 1

To evaluate how temperature and pH affect process stability during continuous production, we conducted iDOE1 using a two-factor, three-level full factorial design. The focus was on identifying process conditions that support stable biomass growth and consistent AA production over time. Process stability was assessed by monitoring the variation in AA composition under different setpoint combinations across several reactor runs. Each condition was maintained for three working volume exchanges (10 d), providing sufficient time for the culture to physiologically acclimate to the new environmental conditions. The measured online values of varied process parameters and their resulting AA composition are shown for R4 over a process duration of 36 d in Figure 2, where circles indicate sampling timepoints. Stable AA production was characterized by the absence of significant changes in the distribution of alanine, glutamic acid and residual AAs in the final section of the stabilization phase of 3 volume exchanges or 10 d. The most stable condition could be identified at pH = 7 and T = 60 °C which corresponds to the process time between 10 and 20 d in Figure 2, approximately, while other conditions clearly do not reach stable production during the 10 d period. However, as visual inspection alone may be misleading or biased by coincidental fluctuations, a quantitative measure was needed to objectively assess process stability across conditions. Therefore, the process stability of each setpoint was further evaluated using the sum of coefficients of variation (CVs), calculated for key outputs including alanine, glutamic acid, total AAs, and biomass X (Supplementary figure 2). All conditions tested in replicate runs exhibited consistent behavior – either relatively stable or unstable – except for setting 5/8 (pH = 6, T = 65O°C) and setting 6/11 (pH = 6, T = 60O°C), which showed divergent outcomes between replicates. This lack of reproducibility suggests that these conditions may lead to inherently unstable and thus unreliable process performance. It should be noted that stability here is assessed relative to the most stable condition observed (pH = 7, T = 60O°C). Given its consistently low variability and robust performance, pH = 7 and T = 60O°C were selected as the standard setpoints for all subsequent continuous experiments at 2OL and 150 L scale.

**Figure 2:**
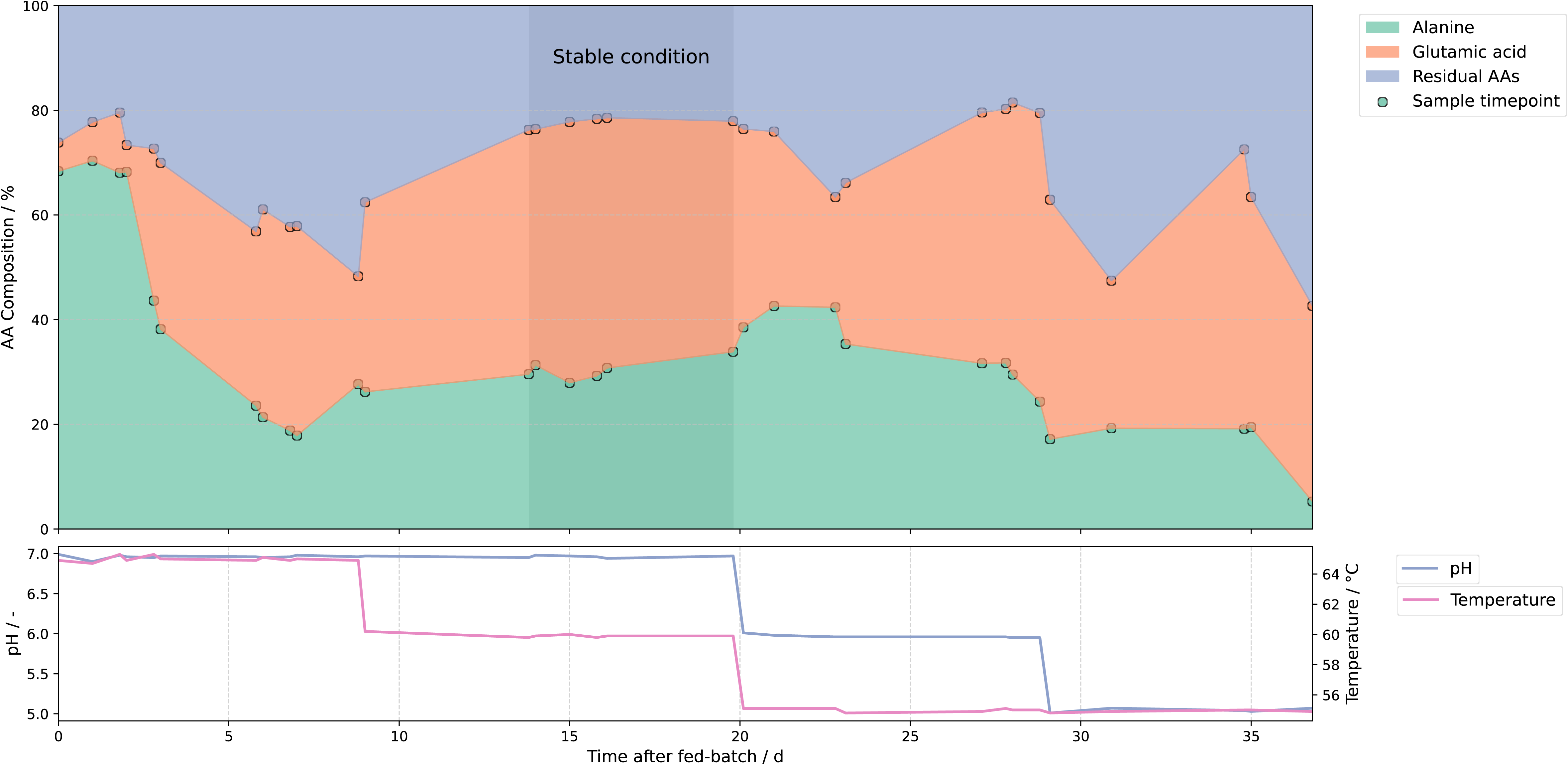
Results of reactor 4 from iDOE1. (A) The relative AA composition over time, expressed as percentage of total AAs, is shown for alanine (green), glutamic acid (orange), and residual AAs (blue). Sample timepoints are indicated by open circles. The shaded region marks a phase of process conditions allowing for stable production, prominent especially between 14 – 20 d which corresponds to 1.5 – 3 volume exchanges at these process settings. (B) Corresponding time-resolved online process parameters: pH (blue) and temperature (pink).

### 2.3. iDOE 2

To further improve process performance following the stabilization using temperature and pH in iDOE1, the second iDOE focused on gas transfer-related parameters. The aim was to optimize Y_AA/x_ by systematically varying stirrer speed, vvm and the H_2_/CO_2_ ratio in the inlet gas. All available data (16 conditions in 8 iDOE runs) contributed to the development of the hybrid model.

To assess the relative influence of the input variables and to evaluate potential redundancy between interdependent parameters such as vvm and k_L_aH_2_, we performed a Sobol global sensitivity analysis. The results show that biomass formation is primarily governed by vvm (S_1_ = 0.26, S_t_ = 0.57) and H_2_ fraction (S_1_ = 0.23, S_t_ = 0.57), both exhibiting strong direct effects and substantial interaction contributions. *γ̇_avg_* also contributes to biomass formation (S_1_ = 0.13, S_t_ = 0.34), albeit to a lesser extent. In contrast, k_L_aH_2_ shows a negligible first-order effect (S_1_ < 0.00) but a pronounced total-order contribution (S_t_ = 0.39), indicating that its influence on biomass arises predominantly through interaction with other variables (Supplementary figures 3 and 4). For total AA formation, k_L_aH_2_ dominates the response (S_1_ = 0.49, S_t_ = 0.70), followed by *γ̇_avg_* (S_1_ = 0.24, S_t_ = 0.34), while vvm and H_2_ fraction exhibit negligible direct effects (S_1_ < 0.00) but non-negligible total-order indices (S_t_ = 0.15 – 0.18), indicating that their contribution arises mainly through interaction (Supplementary figure 5 and Supplementary figure 6). Because the yield Y_AA/x_ is defined as the ratio of total AAs to biomass, these combined sensitivities jointly shape the response. Notably, pronounced total-order and second-order interaction effects, particularly involving vvm and k_L_aH_2_, indicate that the hybrid model captures nonlinear, physically meaningful dependencies. The distinct sensitivity patterns of vvm and k_L_aH_2_ further indicate that these variables are not redundant: while k_L_aH_2_ dominates total AA formation through strong direct effects, vvm contributes primarily via interactions, and conversely exhibits a strong direct influence on biomass formation. This complementary behavior suggests that both variables capture different aspects of the process rather than representing collinear model inputs.

The model was applied within the experimental design and identified an optimal setpoint for maximizing Y_AA/x_ during steady state production at 0.25 L^−1^ min^−1^ vvm, 750 rpm stirrer speed and 90% H_2_ in the gas inflow (corresponding to k_L_aH_2_ = 0.0345 s^−1^ and *γ̇_avg_* = 1162 s^−1^). Although this setpoint was impacted by early termination due to technical issues and laboratory scheduling constraints, it still informed the selection of a targeted validation run. This validation was carried out under both the best-performing iDOE2 conditions and the model-predicted optimum to assess predictive accuracy. The results confirm the hybrid model’s ability to reliably capture the process behavior, particularly during the steady state phase (Figure 3). Under the initial process conditions prior to optimization, an undesired AA profile was produced at an average Y_AA/x_ of 138 mg g^−1^ (±35 mg g^−1^ SD) during steady state (from 7 d onwards). In comparison, the validation run conducted under model-predicted optimal conditions achieved an optimized AA profile, with a 683% increased average Y_AA/x_ of 943 mg g^−1^ (±69 mg g^−1^ SD). The best performing run during the iDOE2 still produced the desired AA pattern, as it was operated using the developed TE solution, however at an average of 617 mg g^−1^ (±85 mg g^−1^ SD) the process conditions only amounted to 447% Y_AA/x_ compared to the initial conditions.

**Figure 3:**
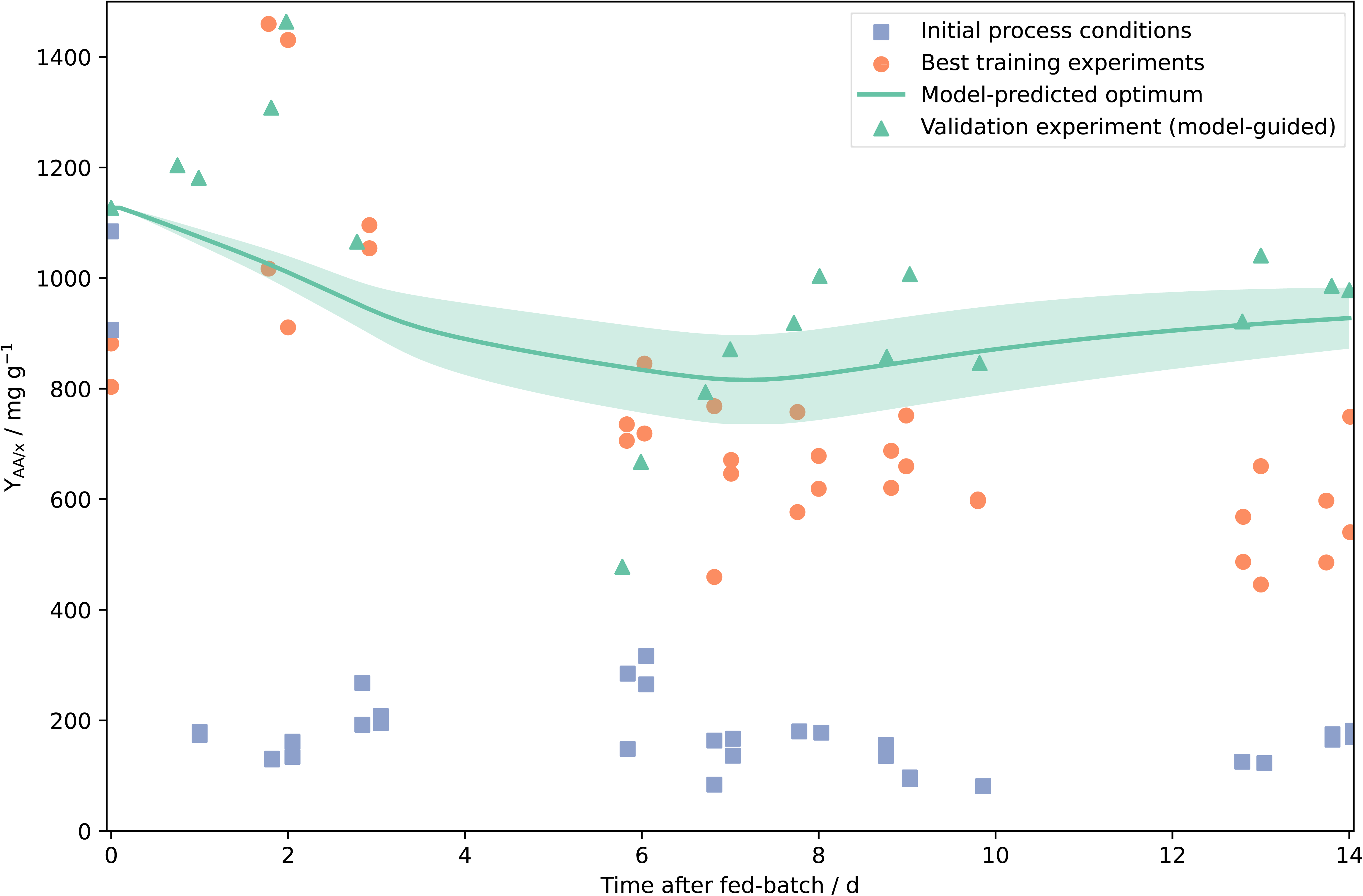
Results of the process optimization using hybrid model predictions for specific biomass yield, trained on iDOE2. Time-resolved measurements of Y_AA/x_ are shown following the transition to continuous cultivation under different operating conditions. Blue squares represent measurements from two independent continuous runs operated under the initial process conditions prior to optimization (vvm = 0.25, agitation = 1500 rpm, pH = 7, T = 65 °C, 80% H_2_ in gas inflow). Orange symbols show measurements from two independent iDOE2 runs that achieved the highest Y_AA/x_ and therefore the best conditions in the model training dataset (vvm = 0.175, agitation = 750 rpm, pH = 7, T = 60 °C, 85% H2 in gas inflow). The green line indicates the hybrid model-predicted optimum (vvm = 0.25, agitation = 750 rpm, pH = 7, T = 60 °C, 90% H_2_ in gas inflow) with the shaded area representing the minimum and maximum predictions across the ensemble models at each time point. Green triangles correspond to an independent validation experiment performed under model-predicted optimal conditions.

### 2.4. Scale-up

Following the identification of optimal process conditions in iDOE2, a scale-up experiment was conducted in a 150 L bioreactor to evaluate whether the optimized AA profile and increased *γ̇_avg_* could be reproduced under technically feasible large-scale conditions while assessing the predictive performance of the hybrid model. Scale-up was performed by selecting operating conditions that reproduce the CFD-derived hydrodynamic descriptors identified at laboratory scale, specifically k_L_aH_2_ and *γ̇_avg_*, within the physical and engineering constraints of the pilot-scale system. This descriptor-based approach aimed to achieve comparable reactor environments relevant to process performance. The scale-up experiment was carried out at T = 60 °C, pH = 7, vvm = 0.18, H_2_ gas inflow = 80 %, stirrer speed = 700 rpm and pressure = 2 bar, which corresponds to k_L_aH_2_ = 0.0168 s^−1^ and *γ̇_avg_* _=_ 2178 s^−1^. This condition if fully covered in the iDOE2 experimental space when considering only scale-independent parameters. Figure 4 visualizes the coverage of the experimental space, with iDOE2 conditions shown in black and the pilot-scale run highlighted in orange. The pilot condition lies within the boundaries of the space defined by scale-independent parameters, illustrating the design’s relevance for scale-up. As shown in Figure 5A, the model accurately predicted process performance during the stable production phase from 7 d onward. With an average Y_AA/x_ of 726 mg g^−1^ (±43 mg g^−1^ SD) the pilot-scale process achieved a 5.3-fold increase compared to initial conditions in 2 L scale. Time-resolved AA analysis (Figure 5B) of the 150 L experiment demonstrates that the significant glutamic acid reduction achieved through media optimization in 120 mL scale could be replicated at 150 L scale, with average concentrations of 6.8% (±0.7% SD) post-adaption. Alanine levels were higher than anticipated at 30% (±0.5% SD), still enabling approximately 63% (± 0.4% SD) of the AA fraction to be directed towards residual AAs. The complete time-resolved AA composition throughout the pilot-scale cultivation is provided in Supplementary figure 7. Notably, the AA pattern stabilized from 7 d onward, while Y_AA/x_ continued to increase until 10 d.

**Figure 4:**
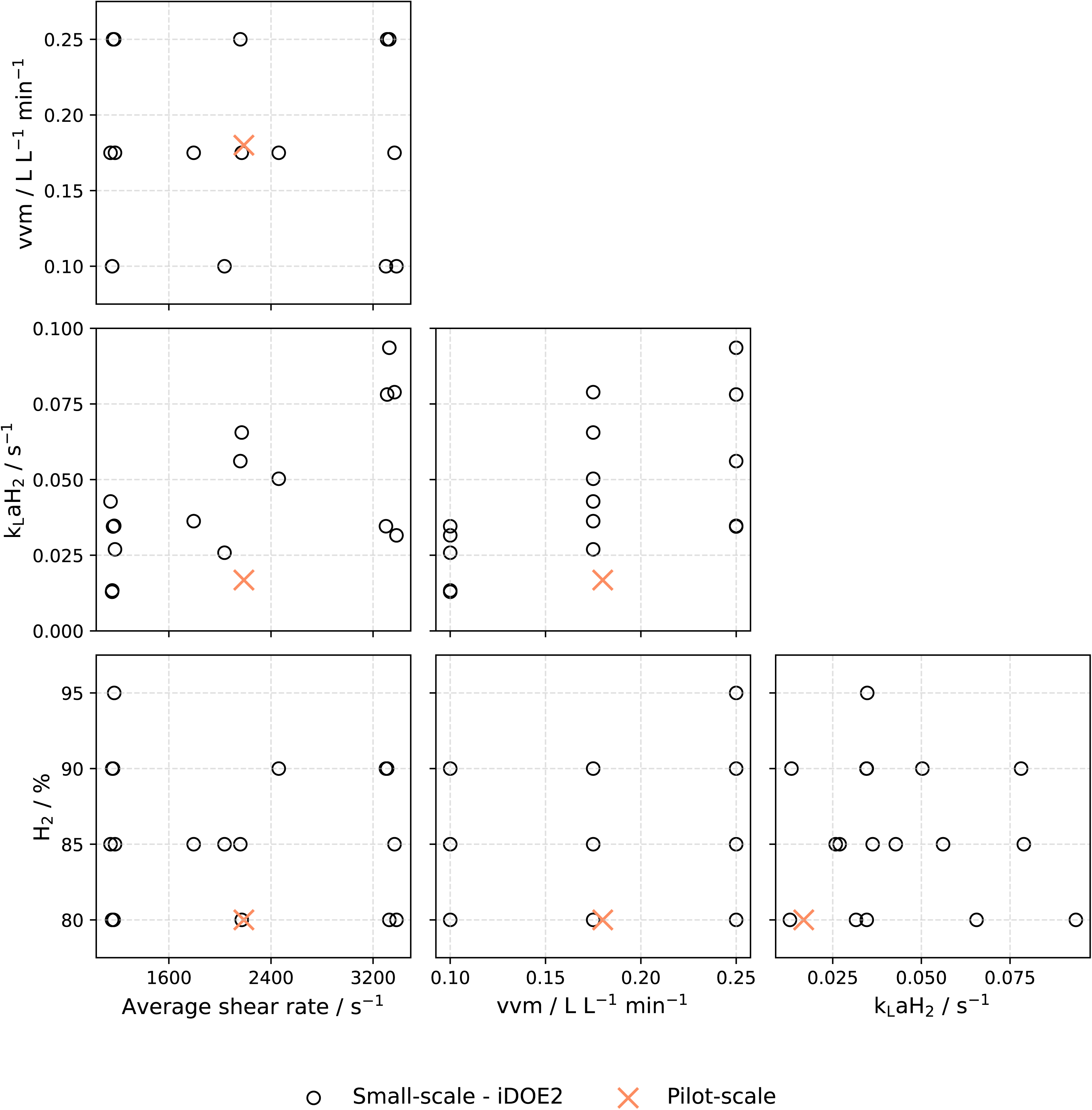
Scatter matrix comparing small-scale and pilot-scale conditions across process parameters covered by the hybrid model. Each subplot shows the pairwise relationship between two input variables, with small-scale points (iDOE2) as empty circles and the pilot-scale condition overlaid as orange cross. The visualization facilitates assessment of the experimental coverage between scales.

**Figure 5:**
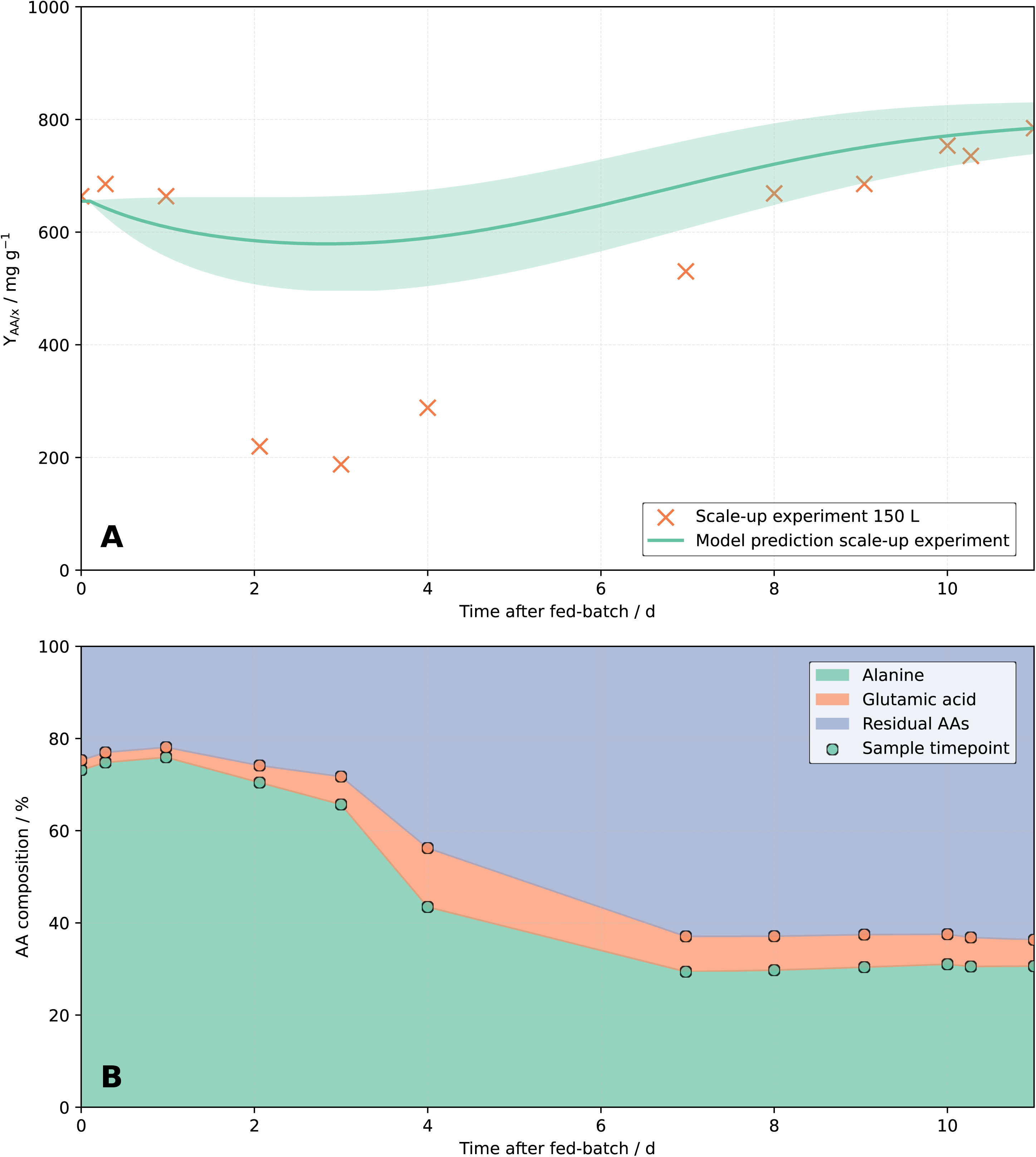
Results of 150 L scale experiment. (A) Y_AA/x_ from offline samples (orange x) and model predictions for 11 d of continuous production, where the shaded region represents the minimum and maximum predictions across the ensemble models at each time point. (B) Measurements of alanine (green), glutamic acid (orange) and residual amino acids (blue) composition for 11 d of continuous production.

### 2.5. Benchmarking of time-resolved and steady-state DOE approach

To benchmark the proposed time-resolved iDOE framework against a conventional steady-state DOE approach, an additional hybrid model was trained using only steady-state observations from the final phase of each experimental condition (last two measurements for x and AAs) while retaining the identical model structure and CFD-derived inputs. The steady-state model identified a substantially different optimum (0.1 vvm, 750 rpm, 80% H_2_, yielding 1254 mg g^−1^) compared to the iDOE model (0.25 vvm, 750 rpm, 90% H_2_, yielding 754 mg g^−1^), demonstrating and overestimation of 66.3% and showing that the two approaches resulted in different process development decisions (Figure 6A). When applied to the pilot-scale process, the steady-state model predicted a Y_AA/x_ of 374 mg g^−1^ after 11 d, whereas the iDOE model accurately predicted the experimentally observed Y_AA/x_ of 785 mg g^−1^ (Figure 6B), which amounts to an underestimation of 52%. Global sensitivity analysis further demonstrated differences in the inferred process relationships. For formation of x, the steady-state model identified k_L_aH_2_ as the dominant direct contributor (S_1_ = 0.35), whereas the iDOE model attributed its influence primarily through interaction effects (S_1_ < 0.00, S_t_ = 0.39, Supplementary figure 8). In contrast, both approaches identified k_L_aH_2_ as the dominant contributor to total AA production, although the steady-state model assigned greater direct contributions to vvm and *γ̇_avg_* (Supplementary figure 9). These results demonstrate that incorporating transient process information improves model identification, resulting in different optimization outcomes and substantially improved predictive performance during scale-up.

**Figure 6:**
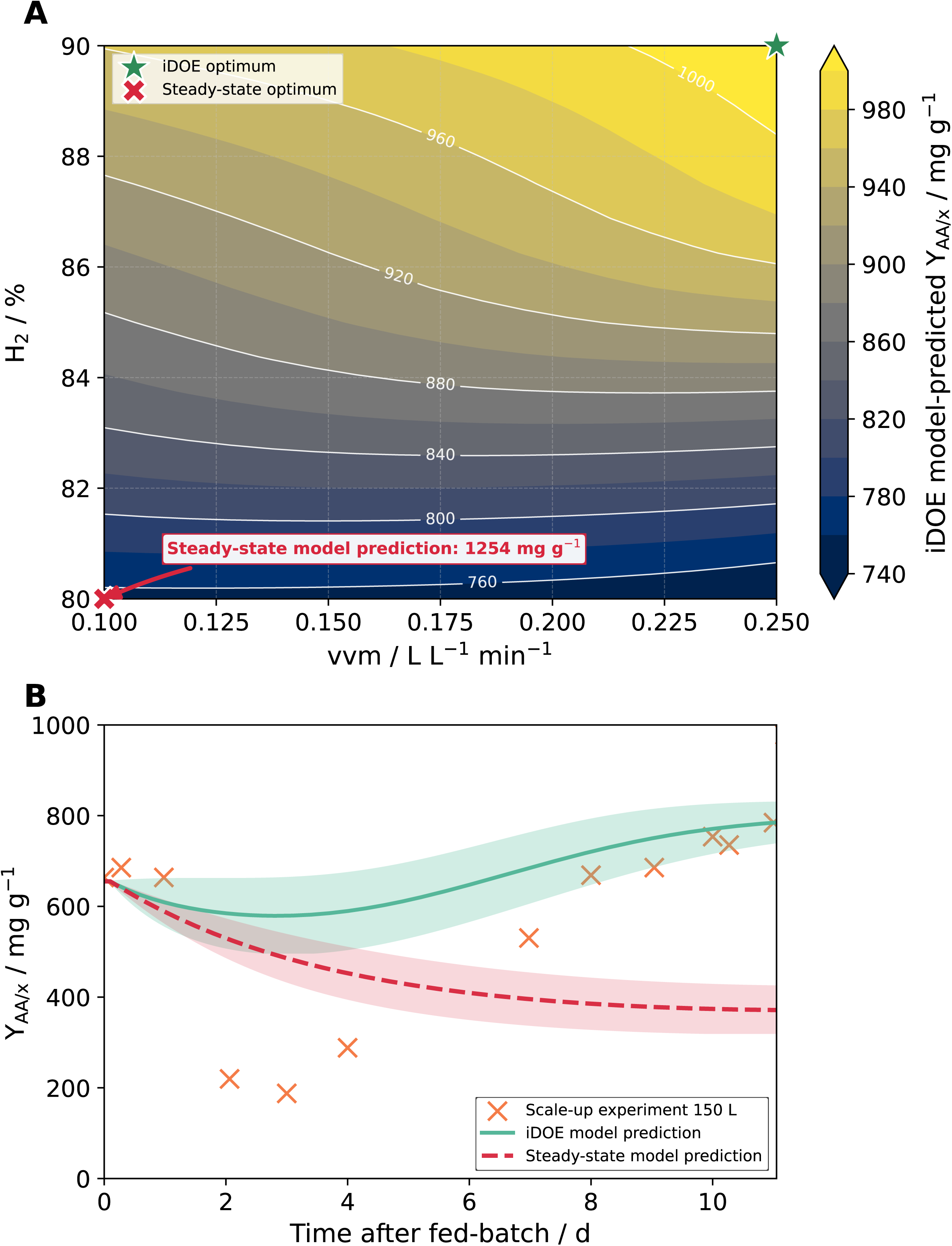
Benchmarking of the time-resolved iDOE framework against a conventional steady-state DOE-approach. (A) Comparison of the optimum operating conditions at laboratory scale predicted by the iDOE model (green star) and a hybrid model trained using only steady-state observations (red cross). Both models employed identical CFD-derived input variables and model structures. The response surface corresponds to the iDOE model prediction at 750 rpm and 14 d of continuous processing, with markers indicating the optimum predicted by each approach. The steady-state model identified a substantially different optimum operating point (red cross, 0.1 vvm, 80% H_2_, yielding 1254 mg g^-1^) compared to the iDOE model (green star, 0.25 vvm, 90% H_2_, yielding 1016 mg g^-1^). The steady-state model overestimated Y_AA/x_ by 66.3% when compared to the iDOE model predictions at 0.1 vvm, 80% H_2_ (754 mg g^-1^). (B) Predictions of Y_AA/x_ during pilot-scale cultivation (150 L). The iDOE model accurately reproduced the yield after 11 d of continuous processing, whereas the steady-state model underestimated process performance at 11 d by 52% (Y_AA/x_ = 374 mg g^-1^). Shaded regions represent the minimum and maximum predictions across the ensemble models at each time point.

## 3. Discussion

Trace elements are indispensable for the function of key metabolic enzymes and play a crucial role in the microbial metabolism, acting as cofactors in enzymatic reactions essential for AA biosynthesis^15,53^. It was shown that *M. marburgensis* is capable of excreting a diverse range of proteinogenic AAs with distinct profiles and that the availability of a nitrogen source affects excretion patterns, particularly for glutamic acid and alanine^32^. We demonstrate that fine-tuning TE composition can control metabolic routing in *M. marburgensis*, shifting away from unwanted AA production with a dominance of alanine and glutamic acid to a redistributed AA spectrum (Figure 1). This expands upon prior findings in *Methanococcus maripaludis*, where Ni and Co supplementation enhanced methanogenic activity and intracellular AA pools^54^. Earlier studies have shown that TE feeding strategies^15^ or supplementation^21,22^ can improve CH_4_ production and influence associated physiological variables. In contrast, the present study focused on redirecting carbon flux towards AA synthesis rather than biomethanation. Thus, formation of CH_4_ was not the main focus of this study and was not analyzed in detail. Our results underscore that TE composition can be deliberately utilized to steer AA biosynthesis towards a targeted pattern, establishing TEs not just as metabolic enablers, but as active levers for bioprocess optimization in methanogens. The observed reduction in total AA concentration under the optimized TE formulation possibly reflects the suppression of energetically favorable overflow pathways, such as alanine and glutamic acid formation, in favor of a more selective and metabolically constrained AA synthesis regime. Mechanistic insights into the regulation of enzymes involved in the AA synthesis pathways leading to a reduction of aforementioned AAs, could not be assessed in the frame of this study. Future work could aim on revealing these regulatory mechanisms on a physiological level by using a combination of omics studies and metabolic modeling.

While different approaches for the optimization of *M. marburgensis* have been reported^55^, the use of iDOE for data generation to describe the steady state performance of a continuous process by leveraging hybrid models is a novelty for this organism. Notably, iDOE significantly reduced experimental time and resource demand to investigate the desired parameter space, as a continuous process no longer needed to be initiated from scratch or run for an extended period of time at constant conditions, thereby enabling faster development cycles.

Beyond the experimental efficiency, process knowledge gathered in this intensified manner is also transferable to larger scales by selecting and scouting the relevant process parameters already early on in process development.

The additional benchmarking against a conventional steady-state DOE approach provides quantitative evidence for the benefit of the proposed time-resolved framework. Although both models employed identical CFD-derived descriptors and model structures, training exclusively on steady-state observations resulted in a different optimum operating point and substantially reduced predictive performance in lab as well as pilot-scale validation experiments. Moreover, differences in the inferred sensitivity indices, particularly for biomass formation, indicate that transient process responses contain information that influence the identification of process relationships beyond steady-state observations alone. These findings suggest that the advantage of the time-resolved approach arises not merely from higher data density, but from capturing the dynamic adaptation of the process following parameter perturbations. Consequently, the resulting hybrid model provides a more reliable basis for optimization and scale transfer while requiring only a single intensified experimental campaign. While the benchmark demonstrates the value of time-resolved data for model identification, appropriate modeling strategies are required to fully exploit this additional information. Fully mechanistic approaches for description of archaeal growth kinetics have been published, however not taking into account the effects of process parameters in a time-dependent manner, nor product formation itself^56^. In contrast, the proposed hybrid model accounts for both time-dependent process conditions and total AA production.

Methanogens depend critically on efficient H_2_ and CO_2_ uptake for energy conservation through hydrogenotrophic methanogenesis, a pathway closely coupled to AA biosynthesis^20,32^. The optimization regarding process performance, focusing on gas-transfer related factors (e.g., k_L_aH_2_, vvm and gassing ratio of H_2_/CO_2_), resulted in an over 6.8-fold improvement of Y_AA/x_ compared to initial performance in 2 L scale, underscoring both the biological relevance of this metabolic link and the predictive capability of the developed hybrid model. Moreover, the gains in Y_AA/x_ offer a strong foundation for further intensification strategies, such as the implementation of cell retention systems^22^, which could decouple biomass accumulation from production and thereby amplify product yields beyond the improvements demonstrated here.

For the first time, we demonstrate that the use of scale-independent process model inputs – supported by rapid-response CFD models and an experimental space feasible across both laboratory and production scale – can enhance process transfer and scale-up. In this way, process knowledge generated during development can be preserved and leveraged to reduce the need for labor- and resource-intensive scale-up experiments. Classical scale-up strategies in upstream bioprocess engineering can for example combine mechanistic kinetic models (e.g., Monod-type formulations) with empirical correlations for gas transfer, in which k_L_a is estimated as a function of the operating conditions. These approaches are widely used and can provide reliable predictions of biomass growth or overall yields, particularly when key parameters are well characterized. However, such correlations are generally system- and geometry-specific and typically describe gas transfer using a single parameter, while additional hydrodynamic factors such as shear or mixing characteristics are not explicitly incorporated despite evidence that such parameters can influence process performance. This has been demonstrated for filamentous microorganisms where the energy dissipation rate was identified as a relevant scale-up parameter and quantified using CFD simulations^57^.

In contrast, the approach presented here combines CFD-derived hydrodynamic descriptors with a hybrid modeling framework, enabling the prediction of more complex outputs such as AA production rates and potentially AA compositions, without requiring an explicit kinetic formulation. The use of CFD-derived descriptors such as k_L_aH_2_ and *γ̇_avg_* enabled the prediction of operating conditions at 150 L that reproduce process-relevant hydrodynamic characteristics observed at small scale. This descriptor-based integration is therefore not intended to replace established mechanistic models, but to complement them in scenarios where multiple interacting process variables and complex product responses need to be captured. Methods in this direction have already been discussed, however by combining mechanistic kinetic models with classical CFD at a transport equation level, that are requiring high computational cost^58^. The approach described above supported robust cell growth and consistent AA production after an initial adaptation phase. Other strategies in this direction have been applied in scaling *M. maripaludis* and *Sulfolobus acidocaldarius*, where physiological markers such as membrane integrity, enzyme activity, and energy yield were prioritized over direct replication of process conditions across scales^59,60^.

In this work, both an increased Y_AA/x_ and an improved AA production profile were successfully translated from 120 mL to 150 L scale, corresponding to a scale-up factor of 2000 when taking into account the working volumes at the respective scales. While pilot-scale Y_AA/x_ matched the predictions of the hybrid model and glutamic acid remained within the expected range, alanine concentrations were elevated compared to the optimized small-scale process, suggesting the influence of additional mechanisms not captured by the tested parameters or TE formulation.

These observations highlight potential areas for improvement and further development. One promising direction is the direct integration of key AAs as output variables in the hybrid model, which could enhance its predictive accuracy – particularly addressing the alanine deviations observed at pilot scale. This extension is currently in development and aims to enable the model to account for AA profile shifts under varying process conditions. In the present work, total AA production and AA composition were predicted using two independent modeling approaches, while integration of these outputs into the hybrid model will support a more unified framework. Moreover, the hybrid model developed in this study is well suited for real-time applications as a digital shadow or even a digital twin, especially when interfaced with appropriate laboratory sensors and equipment. The proposed integration of rapid response CFD simulations with hybrid modeling provides computational times compatible with real-time requirements – an essential feature for dynamic process control^61^. Integrating the AA profile into this dynamic framework would not only enable continuous monitoring but also facilitate advanced control strategies to dynamically steer product composition during continuous operation.

Furthermore, the present hybrid modeling approach relies on spatially averaged CFD-derived descriptors, such as k_L_aH_2_ and *γ̇_avg_*, to enable tractable model identification and scale translation. While these quantities capture key aspects of the overall hydrodynamic and mass transfer regime, they do not explicitly resolve local gradients or zone-specific heterogeneity within the reactor. At larger scales than 150 L, such heterogeneity may become increasingly relevant, particularly in regions near impellers or gas inlets. More detailed representations, such as compartment models or distribution-based descriptors derived from CFD, could provide additional resolution of spatial effects and represent a natural extension of the framework presented here. Incorporating such approaches may further improve predictive accuracy at larger scales, but was beyond the scope of the current study.

While the presented results demonstrate the effectiveness of the hybrid modeling framework for *M. marburgensis* and AA production, the study is currently limited to a single organism and product class. Consequently, the trained model and quantitative results should not be assumed to directly transfer to other archaea or metabolic products. However, the underlying methodology of combining time-resolved experimental design with CFD-informed hybrid modeling provides a general framework that can be adapted to other gas-fermenting microorganisms and production targets, provided that system-specific data are available. Extending this approach to additional organisms and products represents an important direction for future work. The approaches discussed in this study have the potential to considerably reduce time as well as resources spent for optimization in process development, while still gaining the required process insights to scale further.

## 4. Conclusions

This study establishes a robust and scalable strategy for the production of a tailored AA profile using *M. marburgensis*. By combining intensified experimental design methodologies with hybrid modeling, we identified optimal settings for stable AA biosynthesis, enabled the efficient exploration of process conditions and uncovered key TE effects on metabolic routing. The process was successfully optimized toward significantly higher Y_AA/x_, achieving more than a 6.8-fold improvement over initial conditions.

Importantly, we demonstrate the integration of CFD-derived, scale-transferable parameters into the hybrid model, enabling prediction and optimization of Y_AA/x_ across different scales. By combining DOE and modeling techniques, high Y_AA/x_ and an improved AA pattern could be transferred from 120 mL serum bottles via 2 L to a 150 L bioreactor, corresponding to a reaction-volume scale-up factor of 2000. This finding represents a major step forward in gas fermentation technologies and enables AA production from CO_2_/H_2_ by methanogens at larger scales.

Despite common challenges such as gas transfer and mixing limitations at larger scale, the chosen scale-up strategy ensured robust performance after a short adaptation phase. The strong agreement between predicted and observed values for Y_AA/x_ and major AA products supports the reliability of the modeling approaches applied, while remaining deviations (e.g. alanine formation) highlight opportunities for further refinement.

Together these findings advance the design of gas-based bioprocesses using methanogens and establish a broadly applicable framework that complements existing scale-up methodologies, with potential extensions to other cell types, products and industrial sectors.

## 5. Materials and methods

### 5.1. Strains

*Methanothermobacter marburgensis* DSM 2133 was used in all experiments^32,35,62–66^.

### 5.2. Chemicals

H_2_ (99.999 Vol.-%), CO_2_ (99.999 Vol.-%), molecular nitrogen (N_2_) (99.999 Vol.-%), H_2_/CO_2_ (20 Vol.-% CO_2_ in H_2_) (4:1), were used for pre-cultures and in the experimental work. For gas chromatography (GC), CH_4_ (99.995 Vol.-%), molecular oxygen (O_2_) (99.995 Vol.-%), N_2_/CO_2_ (20 Vol.-% CO_2_ in N_2_), H_2_/N_2_ (5 Vol.-% H_2_ in N_2_) plus Argon (99.995 Vol.-%) and Helium (99.995 Vol.-%) were additionally used. All gases were purchased from Air Liquide (Air Liquide GmbH, Schwechat, Austria). All other chemicals were of the highest grade available.

### 5.3. Media

*M. marburgensis* was grown in minimal medium, referred to as MM^53^. To ensure that CO_2_ served as the sole carbon source for both growth and biomethanation, Na_2_CO_3_ was omitted from the MM medium^15,20^. The reduced media recipe can be found in Supplementary table 3. For the purpose of media optimization, the trace element (TE) solution composition was varied in closed batch experiments. Due to intellectual property considerations, the exact TE composition cannot be disclosed in absolute terms. The formulations are therefore reported in relative form where appropriate, while preserving the interpretability of the results^37,38^. Following optimization, the most suitable TE formulation was adopted as the standard solution for further experiments.

### 5.4. Closed batch cultivation

Closed batch cultivation experiments were conducted in 120 mL serum bottles (VWR, Austria). Media was aliquoted to a total working volume of 50 mL and closed with rubber stoppers (20 mm, butyl rubber, CLS-3409-14, Chemglass Life Sciences, USA), pre-boiled 10 times for 30 min in Milli-Q water, and aluminum crimp caps (Ochs Laborbedarf, Bovenden, Germany). An anaerobic headspace was created before autoclavation by gassing with the respective gas mixture H_2_/CO_2_ (4:1) up to 3 bar abs. pressure, repeating the procedure five times^67^. Na_2_S·9H_2_O was added to the sterile media in an anaerobic glove box (Coy Laboratory Products, Grass Lake, USA) prior to inoculation. Sterile syringe filters (w/0.2c µm cellulose, 514-0061, VWR Internation, USA) and sterile needles (disposal hypodermic needle, Gr 14, 0.60 × 30 mm, 23 G × 1 1/4”, RX129.1, Braun, Germany) were used for gassing. Cultivation was performed in a water bath (GFL 1083, Burgwedel, Germany) at 65 °C for 24 h with a start OD of approximately 0.2. To monitor growth and AAs, liquid samples of each 1 mL were withdrawn from the serum bottles. Growth was measured spectrophotometrically via OD (λ = 578 nm, blanked with Milli-Q water) (Hach, DR6000 UV-VIS Spektralphotometer, Vienna, Austria). To quantify the AAs, a liquid sample was taken and centrifuged at 16,100 rcf for 15 min (5415 R, Eppendorf AG, Hamburg, Germany). The supernatant was immediately analyzed. Cell pellets were stored in sterile Eppendorf tubes at −20 °C until further analysis. The relative pressure with the serum bottle was measured using a digital manometer (LEO1-Ei, 1-3 bar rel., Keller GmbH, Winterthur, Switzerland). All measurements were performed at room temperature.

### 5.5. Lab scale continuous culture bioprocess

Continuous cultivation experiments were performed with *M. marburgensis* in a DASGIP® 2.2 L bioreactor system (SR1500ODLS, Eppendorf AG, Hamburg, Germany) with a working volume of 1.6 L, including 100 µL L^−1^ of antifoam (Struktol SB2023, Schill und Seilacher, Hamburg, Germany). Prior to the continuous process, a fed-batch cultivation was performed. The bioreactor was flushed with N_2_ to reach anaerobic conditions followed by H_2_/CO_2_ at a ratio of 4:1 H_2_:CO_2_. CO_2_ was controlled via the MX4/4 unit (Eppendorf AG, Hamburg, Germany) and the H_2_ gas flow was controlled via the C100L Unit (Sierra Instruments, Monterey, USA). Redox potentials and pH values were monitored by individual Redox- and pH-probes (Mettler Toledo GmbH, Wien, Austria). Initial process parameters prior to optimization were temperature of 65 °C, pH of 7 and 1500 rpm, as found suitable for biomass growth in previous studies ^15^. If not indicated otherwise, these set points were used. After anaerobization, 2 mL of 0.5 mol L^−1^ Na_2_S·9H_2_O solution was added to the media. Every fed-batch cultivation was performed with 1% inoculum of a stock culture of *M. marburgensis* that has been adapted through a fed-batch cultivation run before. Feeding of the 0.5 mol L^−1^ Na_2_S·9H_2_O was adapted to the dilution rate of 0.0125 h^−1^ once the fed-batch culture reached an OD of approximately 1, and the continuous phase was started. For media-in and media-out vessels, polypropylene bottles (Nalgene™ Carboy) ranging from 5-20 L were used.

Filter-sterilized RO-water was mixed with autoclaved concentrated media and transferred into the media-in vessels, they were flushed with N_2_ for 30 min, to ensure an anaerobic atmosphere. Liquid samples of 2 mL for AA analysis were taken at each timepoint and centrifuged at full speed (16,100 rcf) for 15 min (5415 R, Eppendorf AG, Hamburg, Germany). Cell pellets and supernatant of each experiment were stored in sterile Eppendorf tubes at −20 °C until further analysis. Gas samples were taken as described below.

### 5.6. Design of experiments

#### 5.6.1. Closed batch design of experiments

Closed batch experiments were conducted to screen for the effects of TE concentrations (CoCl_2_·6H_2_O, FeCl_2_·4H_2_O, NiCl_2_·6H_2_O, MgCl_2_·6H_2_O, Na_2_MoO_4_·2H_2_O) on the resulting AA pattern produced. A total of 60 available serum bottles (section 2.4) were utilized efficiently by combining a fractional factorial screening design with 5 factors and 2 levels, center point runs, negative controls as well as additional runs part of the full factorial design. Each setting was tested in triplicates to ensure reproducibility, while the negative controls were not inoculated. A detailed overview of the experimental design can be found in Supplementary table 4.

#### 5.6.2. Intensified design of experiments 1

The intensified design of experiment 1 (iDOE1) was created to investigate the impact of temperature and pH on overall process stability, which was characterized by a stable growth of biomass, production of AAs and their respective pattern. Ideal utilization of the available laboratory equipment was ensured by splitting up a two-factor, 3-level full factorial design across 4 bioreactors (Supplementary figure 10). The first parameter switch was initiated following an initial stabilization phase of 240 h after switching from fed-batch to continuous processing. The same duration, which is equivalent to 3 working volume exchanges at a dilution rate of 0.0125 h^−1^, was used for each process parameter combination. This duration was determined from preliminary continuous culture runs as a pragmatic balance between experimental throughput and the need to achieve representative steady-state behavior for biomass and AA metrics. The temperature and pH levels were selected around the baseline continuous process used prior to this study and reflect empirically validated operating windows for the strain, ensuring measurable effects on growth and AA production without exceeding known viability or stability limits. Step magnitudes (e.g., ∼5 °C and ∼0.5 pH units) were chosen based on prior experience with the organism to provide sufficient contrast relative to analytical variability. As an additional safety measure and to avoid process failure due to a putative detrimental temperature setpoint and therefore risking a process restart, the iDOE1 was planned to ensure all experiments starting at 65 °C and decrease from there. Several duplicates were included in the experimental design to investigate possible time-dependency of the setpoint combinations. Reproducibility overall was considered in R1 and R4, which share the first 3 setpoints (Supplementary figure 11). For evaluation of stability regarding AA production and growth, the coefficient of variation (CV) was calculated for alanine, glutamic acid, total AA concentration and biomass for each setpoint according to Equation 1. Overall stability was assessed using the sum of CVs for the above mentioned variables per setpoint.

Equation 1: CV using the standard deviation (*σ*) and mean (*μ*).

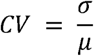

#### 5.6.3. Intensified design of experiments 2

The iDOE2 was developed to focus on gas-transfer related parameters, more specifically the stirrer speed, gas flow coefficient (vvm) and the gassing ratio of H_2_ (in relation to CO_2_) in the gas inflow, and their impact on process performance regarding Y_AA/x_. To utilize the bioreactor capacity of 8 DASGIP vessels ideally, a face-centered central composite design (CCF) was used as basis for the iDOE2. 240 h (3 volume exchanges) were used as stabilization time after switching to continuous procesgising, as well as for each setpoint, as described for iDOE 1. 15 different setpoint combinations and one biological duplicate (centerpoint) could thereby be investigated in 8 vessels over a time period of 30 d. If feasible, experiments were designed to start either at centerpoint or cornerpoint conditions, to minimize the risk of losing essential information regarding process performance by e.g., process failure after a parameter switch. Additionally, physical limitations in laboratory H_2_ supply were considered, as operating multiple high-H_2_-demand experiments concurrently would exceed the facility’s maximum gas flow capacity. The exact sequence of setpoints inside of the design space is shown in more detail in Figure 7. Setpoints for vvm and gassing ratio were defined to span a feasible experimental space for the introduction of changes in gas-liquid transfer and process performance, while remaining compatible with laboratory constraints (e.g., maximum H_2_ delivery) and translatable to pilot-scale operation. The agitation range (750-1500 rpm) was selected to cover the operational window relevant for continuous operation in the 2 L system. Lower agitation rates resulted in insufficient gas–liquid mass transfer, while higher rates approached mechanical limits and did not provide additional benefit. The selected range therefore ensured stable operation while spanning a broad range of hydrodynamic conditions for the DOE.

**Figure 7:**
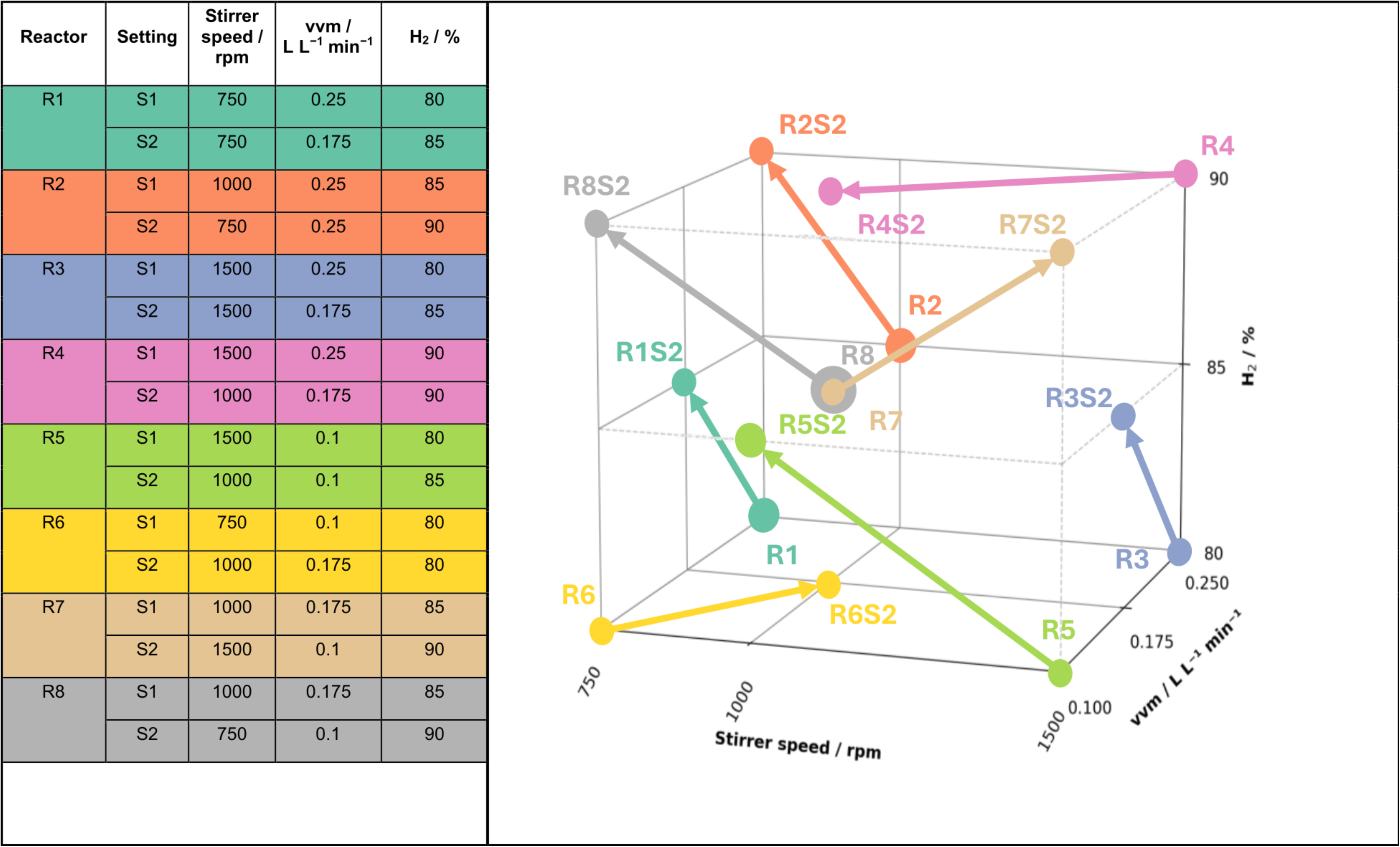
iDOE 2 settings. Each reactor (R1–R8) was operated with dynamic shifts in stirrer speed, vvm and gassing ratio H_2_ setpoints over the course of a single experiment. The left table lists all applied setpoints and their allocation. Each run included one parameter combination shift, as indicated by e.g. R1 (Reactor 1 – setpoint 1) and R1S2 (Reactor 1 - setpoint 2).

### 5.7. Gas chromatography and analysis

Off-gas sampling was performed by flushing a serum bottle, closed with blue rubber stoppers (20 mm, butyl rubber, CLS-3409-14, Chemglass Life Sciences), for approximately 10 min directly connected to the off-gas stream of the bioreactor. The purpose was to ensure that the atmosphere inside the bottle is displaying the actual off-gas in the bioreactor. Gas samples were taken after 3x volume exchange, always in duplicates. The off-gas composition (H_2_, CO_2_, CH_4_ and N_2_) of the collected gas samples were analyzed using the Teckso Gas Chromatograph (Agilent 990 Micro GC, Agilent Technologies, Santa Clara, CA, USA) with a thermal conductivity detector (TDC) and two independently controlled channels. Channel 1 uses argon as carrier gas and is used for measuring H_2_, N_2_, O_2_ and CH_4_, while CO_2_ is measured within channel 2 using helium as carrier gas. The injection temperature for both channels was set to 110 °C, column temperature was 60 °C for channel 1 and 70 °C for channel 2 and the detector pressure was set to 150 kPa. A total of 16 gas samples could be serially connected to the system via needles and tubing. As first measurements often showed traces of O_2_ that was still trapped in the sampling lines, therefore only second measurements were taken for further analysis. The results were then normalized to 100% and averaged per duplicate. Using the same method, the machine was periodically calibrated with a minimum of 8 different standard gases (100% H_2_, 100% O_2_, 100% N_2_, 100% CH_4_, 100% CO_2_, gas mix of 80% H_2_ and 20% CO_2_, a gas mix of 20% CO_2_ in N_2_, a gas mix of 5% H_2_ in N_2_).

### 5.8. Liquid sample analysis

Microbial growth was measured spectrophotometrically via OD (λ = 578 nm, blanked with Milli-Q water) (Hach, DR6000 UV-VIS Spektralphotometer, Vienna, Austria). NH_4_^+^ determination was performed with the same device and the ammonium kit (Hach, LCK303 and LCK502, Vienna, Austria).

### 5.9. Amino acid analysis

AA analysis (20 proteinogenic including homoserine, hydroxyproline, norvaline, taurine and ornithine) were quantified directly from growth medium using a diluted, cell-free supernatant. It was performed by HILIC-LC-MS/MS with ESI and external calibration as reported before^35^. Selected figures in the main manuscript display grouped AA compositions, whereas complete AA profiles are provided in the supplementary information (Supplementary figure 1, Supplementary figure 7, Supplementary table 2).

### 5.10. Modeling

#### 5.10.1. Media optimization

Media optimization was performed based on the results from the closed batch DOE. Second-degree polynomial regression models were developed for each response variable (alanine (%), glutamic acid (%), and total AA concentration (mg L^−1^)), using the PolynomialFeatures and regression functions from the scikit-learn Python library^68^. The predictive models were integrated into a single scalar objective function to enable multi-objective optimization using the Sequential Least Squares Programming (SLSQP) algorithm implemented scikit-optimize. To ensure comparability between response variables with different units and numerical ranges, all model outputs were normalized to a dimensionless range of [0,1] using min–max scaling. The normalization bounds were defined based on the minimum and maximum values observed within the DOE experimental space and applied to model predictions during optimization.

Equation 2: Objective function for optimization using SLSQP algorithm.

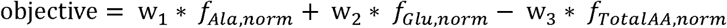

*f_i_*_,*norm*_ denotes the normalized response variables and the maximization of total AA concentration was incorporated into the minimization framework by subtracting the corresponding normalized term.

The weighting factors (w_1_=1.2, w_2_=2.0, w_3_=0.5) were selected to reflect the relative optimization priorities. In particular, glutamic acid reduction was assigned the highest weight due to its dominance in the amino acid spectrum and its strong impact on the desired product profile. Alanine reduction was assigned a secondary weight, while total AA concentration was included as a supporting objective to avoid solutions with low overall productivity. Due to normalization of all response variables, the weights directly represent the relative importance of each objective. The selected values were chosen to enforce a clear prioritization (Glutamic acid > Alanine > total AAs) while maintaining a balanced trade-off between composition and productivity. Bound and inequality constraints were included to maintain process feasibility and enforce biological limits. Specifically, alanine and glutamic acid concentrations were constrained to remain below 35% and 15%, respectively, while all input variables (TE concentrations) were restricted to the ranges defined by the closed batch DOE experimental space.

#### 5.10.2. Rapid response CFD model

To enable the utilization of CFD derived parameters like volumetric mass transfer coefficient and the average shear rate, so-called rapid response models were created for the laboratory scale reactor described in section 2.5. CFD simulations were used to generate a consistent dataset for the outputs of interest (*γ̇_avg_* and k_L_aH_2_) covering the operating domain of the reactor defined in the DOE. The varied operating parameters for the CFD simulations consisted of stirrer speed, vvm and gassing ratio of H_2_/CO_2_. The resulting data was then used to train a regression model capable of predicting named outputs across the entire operating range of the utilized bioreactor. Once trained, the rapid response model provided instantaneous predictions of time-intensive numerical simulations at any arbitrary combination of inputs capturing the underlying trends of the outputs of interest.

Consequently, real-time reactor operating parameters could be utilized to compute scale-independent *γ̇_avg_* and k_L_aH_2_ values, which were subsequently used as inputs for training the hybrid model.

##### 5.10.2.1. Numerical methodology

CFD simulations were performed using the commercially available simulation software developed by SimVantage (SimVantage GmbH, Graz, Austria, available at: https://simvantage.com/features/)^57^, which is specifically tailored to the use case of stirred tank bioreactors. The solver is based on the lattice-Boltzmann method^69^ for incompressible fluid flow. Gas-liquid bubbly flow was modeled with a two-way coupled Eulerian-Lagrangian approach, where the liquid phase was resolved on the Eulerian lattice and the dispersed gas phase was represented by Lagrangian bubbles. Turbulence is considered by means of a large eddy simulation (LES), with subgrid-scale effects being modeled with the Smagorinsky-Lilly model. Interfacial mass transfer was modeled with the Eddy-Cell model by Lamont and Scott^70^, which relates the local mass transfer coefficient to the turbulent energy dissipation rate in the liquid at the bubble interface. The average shear rate was computed as a volume-averaged quantity over the liquid phase. Detailed descriptions of the models utilized by the software and their validation are provided elsewhere^71–73^.

##### 5.10.2.2. CFD model and operating domain

The implemented geometry of a single DASGIP® reactor is displayed in Supplementary figure 12 with visualizations of the Lagrangian bubbles as well as the velocity field. To derive the selected outputs (*γ̇_avg_* and k_L_aH_2_) from the CFD model over the operating domain defined by the iDOE2, a total of 108 simulations were performed using a structured parameter grid comprising six stirrer speed levels, six gas flow rate (vvm) levels, and three inlet gas composition (H_₂_/CO_₂_) levels. Fluid properties were assumed to be constant over the operating domain and set to 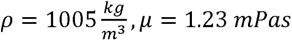 and 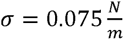.

All simulations were carried out on a uniform Cartesian grid spanning 80 computational nodes across the reactor diameter. This grid resolution was selected as a deliberate compromise between numerical fidelity and computational efficiency, ensuring a reasonable balance between result accuracy and computational feasibility for the simulation of 108 operating conditions across the investigated domain. To assess whether the chosen resolution adequately captures the dominant impeller-driven energy input, the power number *N_P_* was evaluated based on impeller torque and the volume integrated energy dissipation rate. For the employed three-stage Rushton turbine configuration, the resulting power numbers were approximately constant over the investigated Reynolds number range (*Re* ≈ 2 x 10^44^ - 4 .x 104) reflecting the expected behavior in a fully turbulent regime ^74^. As shown in Supplementary figure 13, the magnitude of the computed power numbers and of the respective Reynolds numbers are consistent with literature values ^75,76^ reported for Rushton turbines operating under turbulent conditions. The difference between torque- and dissipation-based power input, lying within 15% over all investigated impeller speeds, is consistent with LES-based stirred tank simulations reported in the literature and reflects expected under-resolution of dissipation at the smallest scales^77^. This level of agreement is considered sufficient to capture the quantities of interest while remaining within a computationally feasibly timeframe for the amount of conducted simulations.

##### 5.10.2.3. Machine learning based rapid response model

CFD results were used to train a machine learning model with Python based on a Kriging partial least squares approach (KPLS)^78^. Parity plots of predictions vs. CFD data for *γ̇_avg_* and k_L_aH_2_ are shown in Supplementary figure 14. The plots indicate that the trained models reproduce the CFD response surfaces with sufficient accuracy across the investigated operating domain and are suitable as rapid response models for the hybrid modeling workflow.

#### 5.10.3. Hybrid model

Machine learning was used in combination with a set of mechanistic equations describing cell growth and productivity to ultimately predict Y_AA/x_ in the continuous phase of the bioprocess. The hybrid modeling framework was implemented using the Novasign Hybrid Modeling Toolbox (Novasign GmbH, Vienna, Austria, version 3.26, available at: https://novasign.at/products/hybrid-modeling-toolbox/)^42^.

##### 5.10.3.1. Datasets

The training/validation dataset included time-resolved information about online and offline data from 8 runs (iDOE 2). For each CPP combination, a new run id was created in the dataset, resulting in 16 individual runs. Utilizing the rapid response CFD model, the two parameters k_L_aH_2_ and *γ̇_avg_* were included in the dataset for each timepoint, depending on the corresponding online values of vvm, stirrer speed and ratio of H_2_ in the gas inflow. The dataset for testing the scale-up model was retrieved from a scale-up run at 150 L and pre-processed in the same manner.

##### 5.10.3.2. Model structure

The model inputs were selected to allow for prediction across different process scales, by substituting stirrer speed with scale-independent parameters k_L_aH_2_ and *γ̇_avg_* in addition to the remaining iDOE factors, being vvm and the ratio of H_2_ in gas inflow. For capturing process performance, an artificial neural network (ANN) was used to estimate specific rates of the respective response variables in a mechanistic framework, being the specific growth rate µ / h^−1^ and the associated production of AAs per biomass q_p_ / mg g^−1^ h^−1^ for the prediction of two outputs, biomass concentration (x / g L^−1^) and total AA concentration (A_t_ / g L^−1^) and the final derived output Y_AA/x_ (mg g^−1^). The ANN was configured with 1 hidden layer encompassing 12 hidden neurons. The hyperbolic tangent (tanh) activation function was used for the hidden layer and z-score scaling was applied to all inputs. The hybrid model structure is shown in more detail in Figure 8.

**Figure 8:**
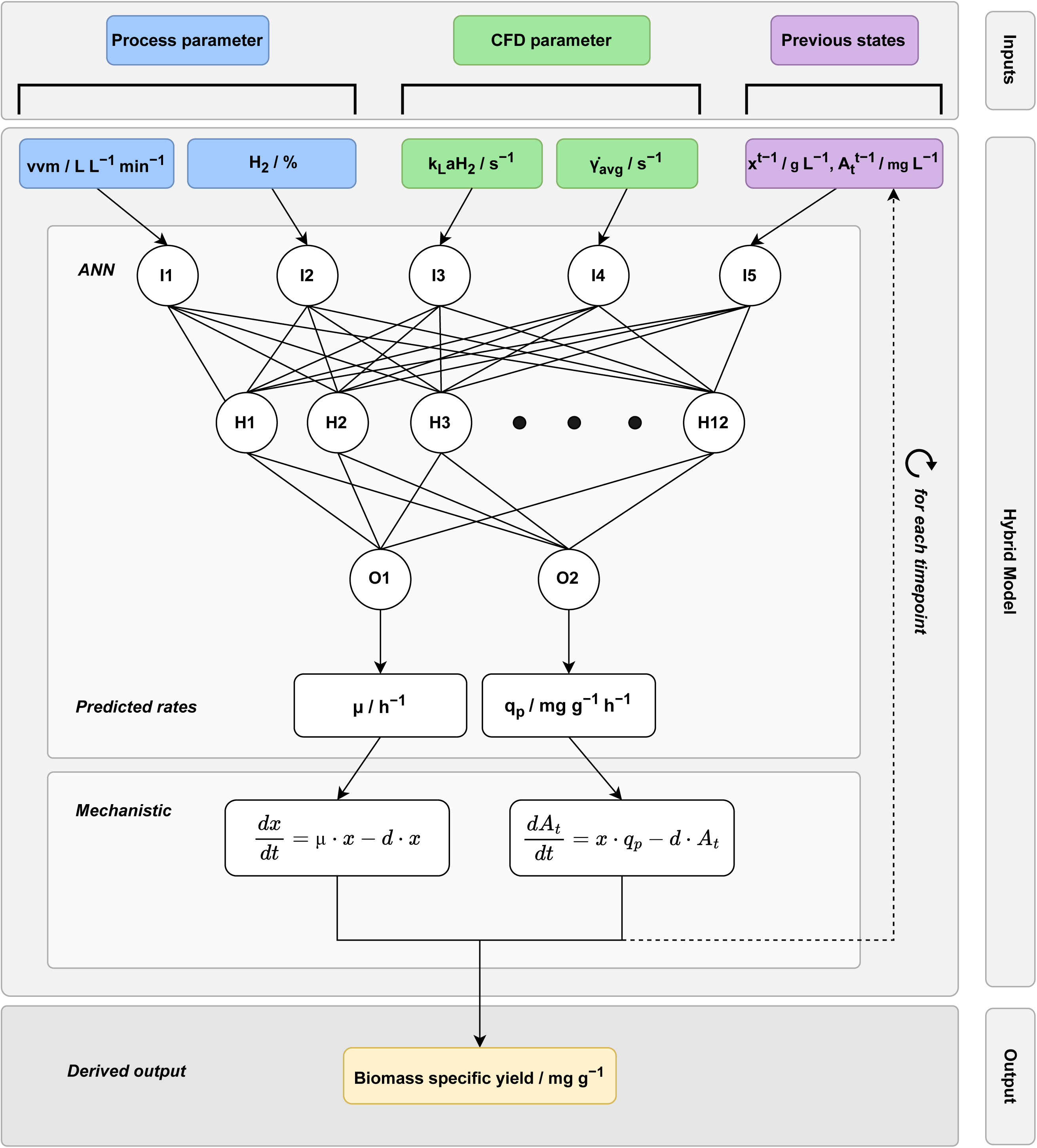
Structure of the hybrid modeling framework integrating process parameters, CFD outputs and previous process states. The framework consisting of an ANN and a mechanistic model is used to predict microbial growth and product formation in the gas fermentation process. The ANN was set up to have 5 input nodes (I1 – I5), receiving two process parameters (vvm, gassing ratio), two CFD-derived parameters (k_L_aH_2_ and *γ̇_avg_*), and the previous states of the biomass concentration (x^t−1^) and total AA concentration (A_t_^t−1^). Via 12 hidden neurons (H), the ANN predicts the two outputs specific growth rate (μ) and specific productivity (q_p_) (O1 and O2), which serve as input rates for a set of ordinary differential equations (ODEs) describing biomass (x) and total AA accumulation (A_t_) over time. These ODEs include a dilution term (d), and the ANN predictions are recalculated for each timepoint using the previous predictions as inputs (dashed line). From the resulting trajectories, Y_AA/x_ is derived for each timepoint as the final model output. Colored labels denote the origin of the respective input parameters (blue: process parameters; green: CFD outputs; purple: previous states), while grey boxes group related components of the hybrid framework.

##### 5.10.3.3. Hybrid model training and evaluation

The training dataset comprising 16 runs was split into training (15 runs) and validation (1 run) sets in a run-wise manner, and this procedure was repeated eight times (eight bootstrap iterations). Model training was initialized with 16 parallel starts to increase the likelihood of convergence to a global rather than a local minimum. The Levenberg–Marquardt optimizer was reset four times, every 50 iterations, during the training process. The optimization minimizes a least-squares objective between model predictions and measured data, with step sizes and convergence handled internally by the Levenberg–Marquardt algorithm. The model selection and evaluation was based on the normalized root mean square error (NRSME) using Equation 3, with the number of datapoints (*n*), observed values (*y_i_*), predicted values (*ŷ*) and the mean of observed values (*ȳ*). An ensemble model was created using 3 models from different boots to avoid overfitting but still include all information available.

Equation 3: NRMSE calculation

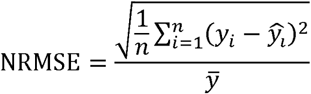

#### 5.10.4. Sensitivity analysis

A global variance-based sensitivity analysis was performed using the Sobol method implemented in the SALib Python package (SALib.sample.sobol for sampling (n=512) and SALib.analyze.sobol for index calculation)^79^. First-order (S_1_), total-order (S_t_), and second-order interaction (S_2_) Sobol indices were computed for all model inputs to quantify their individual and joint contributions to the variance in biomass and total AA concentration. Input bounds were defined consistently with the experimental space.

#### 5.10.5. Process simulation

Utilizing the Novasign Digital Twin (Novasign GmbH, Vienna, Austria, version 1.5.0, available at: https://novasign.at/products/digital-twin/), a simulation comprising the parameter space shown in Figure 7 was created, which was enabled by calling the rapid response CFD model for each simulated point in the 3-dimensional space, thereby establishing a quantitative relationship between stirrer speed, k_L_aH_2_, and *γ̇_avg_* . Simulation results further allowed for the model-based prediction of a process optimum regarding Y_AA/x_.

### 5.11. Scale-up of the continuous culture bioprocess

The continuous fermentation was scaled up to a 150 L continuous stirred-tank reactor (CSTR) (customized pilot fermenter P M0 150 L from Bioengineering AG, Switzerland). The bioreactor was filled with 96.5 L RO water including salts (final concentrations in 100 L: 2.1 g L^−1^ NH_4_Cl, 2.62 g L^−1^ KH_2_PO_4_, 5.36 g L^−1^ K_2_HPO_4_), anaerobized with N_2_ and sterilized in place at 116 °C for 40 min. The vacuum was broken with N_2_ gas to avoid introduction of O_2_. An O_2_ probe monitored anaerobic conditions inside the bioreactor. After cooling down to 60 °C, H_2_/CO_2_ (4:1) gassing was started at 0.1 vvm as well as pH control with 2 mol L^−1^ NaOH as basic agent to counteract acidification through CO_2_ gassing. The stirrer was commenced at 200 rpm. 100 mL of autoclaved optimized TE solution and 400 mL of 0.33 mol L^−1^ Na_2_S·9H_2_O solution were added via feed pumps. The AwiFLEX XL gas analyzer (AWITE Bioenergie GmbH, Germany) was started to monitor H_2_, CO_2_, CH_4_ and hydrogen sulfide concentrations in the off-gas. The stirrer speed was increased to 495 rpm while the pressure remained at 1 bar. A sample was withdrawn before and after 500 mL of inoculum of *M. marburgensis* was added via a feed pump to reach an OD of 0.1. The process was operated in fed-batch mode until growth of the organism reached an OD of approximately 1 (3 d), at which point the liquid outflow and therefore the continuous process was started. Fresh medium with the same composition as for the fed-batch mode, mixed in a tank (200 L, PE with stirrer, Schwarzer Rührtechnik GmbH, Germany), UV sterilized (PURION 500, UV Concept GmbH, Germany) and provided in a 300 L buffer tank (MDPE, Tanks Direkt GmbH, Germany), was fed at a rate of 0.0125 h^−1^ into the bioreactor. Na_2_S and the optimized TE solution were supplied separately to avoid the precipitation of salts. A weight-based control loop maintained the bioreactor level at 100Okg, utilizing the internal scale and associated pump system. The liquid harvest was collected in a separate tank (200 L, PE with stirrer, Schwarzer Rührtechnik GmbH, Germany). Antifoam (Struktol SB2023, diluted 1:10) was added on demand during the fermentation. Liquid samples (10 mL) were withdrawn daily to determine OD, NH_4_^+^, and AA composition. Reproducing comparable hydrodynamic conditions in the continuous phase at 150 L scale relative to the 2 L vessel was chosen as scale-up criterion, enabling hybrid model predictions without extrapolation of the parameter space. For this, CFD simulations were used to guide the scale-up by evaluating the relationship between stirrer speed, gassing rate, and resulting k_L_aH_2_ and *γ̇_avg_*. This enabled the identification of operating conditions that reproduce the CFD-derived hydrodynamic descriptors from small scale as closely as possible within technical constraints, while acknowledging that these descriptors represent effective process characteristics rather than complete equivalence of intrinsic gas transfer rates.

### 5.12. Statistics and reproducibility

The study comprises closed batch screening experiments, laboratory-scale continuous cultivations using intensified design of experiments (iDOE), and pilot-scale validation. Closed batch screening was performed using a fractional factorial design with five factors and two levels, complemented by center points, negative controls, and selected runs of the corresponding full factorial design. Each experimental condition was tested in biological triplicates (n = 3), while negative controls were not inoculated.

Laboratory-scale continuous experiments were conducted using two different designs. iDOE1 followed a two-factor, three-level full factorial design to assess the effects of temperature and pH on process stability under continuous operation. iDOE2 was based on a face-centered central composite design (CCF) to investigate the impact of gas-transfer-related parameters (stirrer speed, gas flow coefficient (vvm) and the gassing ratio of H_2_ (in relation to CO_2_) in the gas inflow) on process performance. In both designs, setpoints were applied sequentially under time-resolved, transient conditions, with stabilization periods of three working volume exchanges. Samples collected over time within a single cultivation represent repeated measurements and were not treated as independent biological replicates.

Continuous laboratory-scale cultivations were reproduced across multiple independent runs to assess stability and model robustness. Pilot-scale validation was performed at 150 L scale using a single biological run (n = 1) at one defined operating setpoint, reflecting the practical constraints of large-scale experimentation.

Descriptive statistics are reported as mean ± standard deviation. Statistical significance was assessed using two-sided Student’s t-tests where explicitly stated. For steady-state comparisons within a single continuous cultivation, statistical tests were applied to repeated measurements taken at distinct time points during steady-state operation and are therefore interpreted descriptively, acknowledging the lack of independent biological replication.

Reproducibility and robustness of the CFD and hybrid modeling workflows were assessed through deterministic CFD simulations over a defined operating domain and through bootstrap-based training and ensemble modeling of the hybrid model.

## Author contributions

BR, EH, AR and HP performed experiments in small-scale and laboratory analysis. FS, MS and SD conducted the 150 L reactor experiment. BR consolidated data. BH performed the data analysis, preprocessing and developed all computational models. BH and MK designed the experiments. LG created the CFD model and rapid response model. BH, BR, FS, MD, SK-MRR and LG contributed to manuscript writing. SK-MRR conceived, designed and supervised research. GB and SK-MRR acquired funding.

## Data availability

The datasets generated and analyzed during this study, including those used for model development, are available in the PHAIDRA repository of the University of Vienna^80^.

## Code availability

The CFD simulations and hybrid modeling analyses in this study were performed using licensed commercial software (commercial CFD software provided by SimVantage (https://simvantage.com/features/) and the Novasign Hybrid Modeling Toolbox (https://novasign.at/products/hybrid-modeling-toolbox/, version 3.26.0)). Due to licensing restrictions, the underlying source code cannot be publicly shared. All experimental datasets used for model development and validation are publicly available, and the model structure, input variables, training procedure, and configuration details are described in the methods. Trained model artifacts and configuration files can be made available upon reasonable request and subject to licensing agreements.

## Acknowledgements

We thank Bart Akerboom for his support during cultivations and Nino Trattnig for performing AA analyses.

