## Supplementary information for "CFD-Informed Hybrid Modeling Unlocks Scalable, Tunable Amino Acid Production in *Methanothermobacter marburgensis*"

Austria

Supplementary table 1**: Results of media optimization indicating predicted optimum for 5 trace elements to decrease alanine and glutamic acid production**

| Trace element salt | Level |
| --- | --- |
| CoCl_2_·6H_2_O | − |
| FeCl_2_·4H_2_O | 0 |
| NiCl_2_·6H_2_O | + |
| Na_2_MoO_4_·2H_2_O | − |
| MgCl_2_·6H_2_O | − |

*The model suggested decreasing (“−“), increasing (“+”) or the keeping the concentration (“0”) the respective salt concentrations in comparison to the original media composition. The exact trace element composition is part of an undisclosed process and therefore cannot be reported in absolute terms.

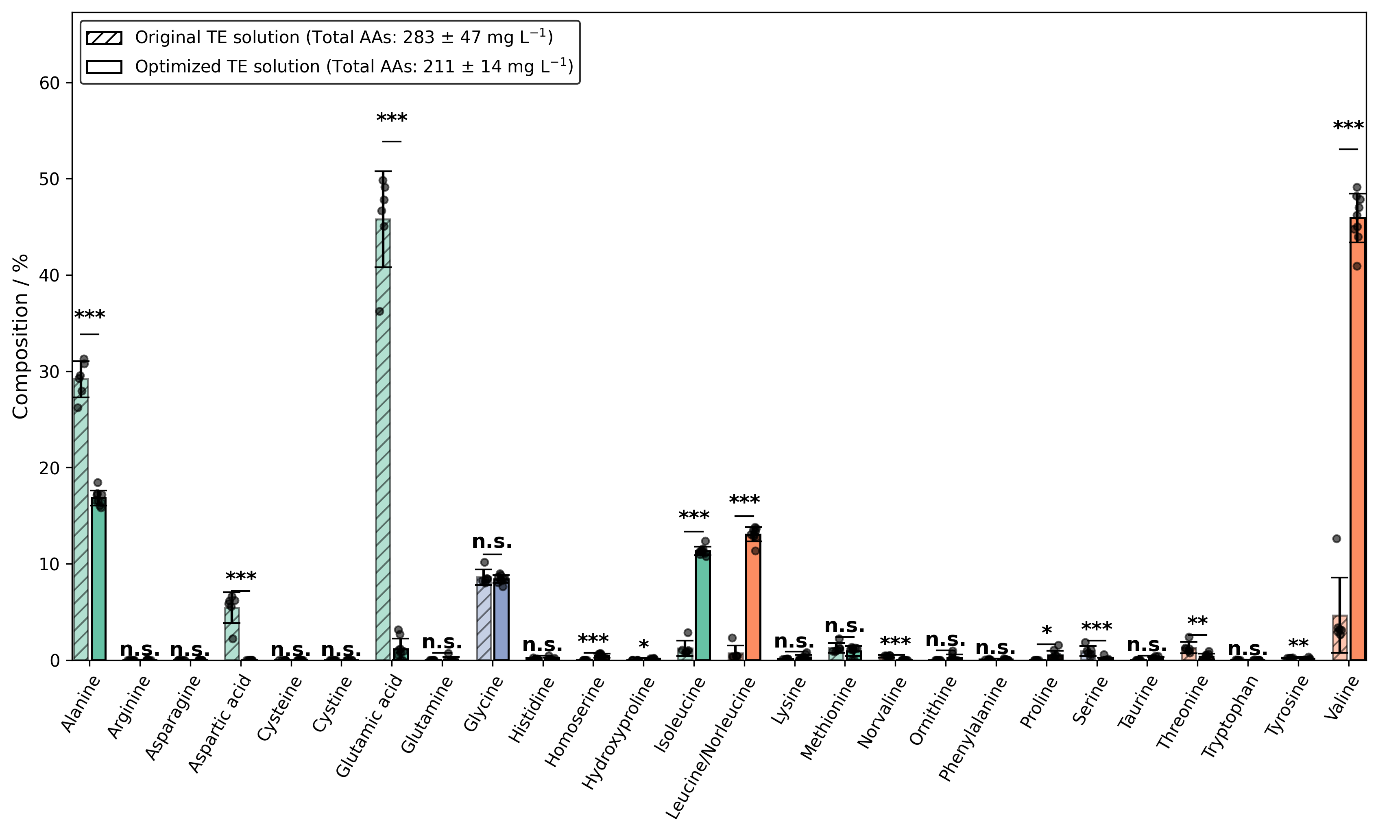

Supplementary figure 1: **Complete amino acid composition before and after trace element optimization.** Bars indicate the mean relative composition of amino acids (AAs), calculated from time-resolved samples collected during the steady-state phase. Individual points represent all time-resolved steady-state measurements. Error bars denote the standard deviation of these measurements within each condition. Statistical significance between original and optimized TE solutions was assessed using two-sided Student’s t-tests on time-resolved steady-state samples (*** p < 0.001, ** p < 0.01, * p < 0.05, n.s. not significant).

| Amino Acid | mean before % | sd before % | n before | mean after % | sd after % | n after | diff % | t | p | Sig |
| --- | --- | --- | --- | --- | --- | --- | --- | --- | --- | --- |
| Alanine | 29.18 | 1.88 | 6.00 | 16.83 | 0.79 | 9.00 | -12.34 | 15.25 | 0.00 | *** |
| Arginine | 0.00 | 0.00 | 6.00 | 0.00 | 0.01 | 9.00 | 0.00 | -1.00 | 0.35 | n.s. |
| Asparagine | 0.00 | 0.00 | 6.00 | 0.00 | 0.00 | 9.00 | 0.00 |  |  | n.s. |
| Aspartic acid | 5.45 | 1.61 | 6.00 | 0.00 | 0.00 | 9.00 | -5.45 | 8.29 | 0.00 | *** |
| Cysteine | 0.00 | 0.00 | 6.00 | 0.00 | 0.00 | 9.00 | 0.00 |  |  | n.s. |
| Cystine | 0.00 | 0.00 | 6.00 | 0.00 | 0.00 | 9.00 | 0.00 |  |  | n.s. |
| Glutamic acid | 45.79 | 4.99 | 6.00 | 1.15 | 1.10 | 9.00 | -44.65 | 21.56 | 0.00 | *** |
| Glutamine | 0.00 | 0.00 | 6.00 | 0.08 | 0.24 | 9.00 | 0.08 | -1.00 | 0.35 | n.s. |
| Glycine | 8.61 | 0.79 | 6.00 | 8.41 | 0.40 | 9.00 | -0.20 | 0.56 | 0.59 | n.s. |
| Histidine | 0.12 | 0.07 | 6.00 | 0.07 | 0.15 | 9.00 | -0.05 | 0.76 | 0.46 | n.s. |
| Homoserine | 0.00 | 0.00 | 6.00 | 0.45 | 0.20 | 9.00 | 0.45 | -6.92 | 0.00 | *** |
| Hydroxyproline | 0.00 | 0.00 | 6.00 | 0.08 | 0.08 | 9.00 | 0.08 | -2.92 | 0.02 | * |
| Isoleucine | 1.24 | 0.80 | 6.00 | 11.33 | 0.45 | 9.00 | 10.10 | -28.02 | 0.00 | *** |
| Leucine/Norleucine | 0.72 | 0.77 | 6.00 | 13.06 | 0.74 | 9.00 | 12.34 | -30.84 | 0.00 | *** |
| Lysine | 0.10 | 0.03 | 6.00 | 0.31 | 0.24 | 9.00 | 0.21 | -2.59 | 0.03 | * |
| Methionine | 1.26 | 0.49 | 6.00 | 0.93 | 0.53 | 9.00 | -0.33 | 1.23 | 0.24 | n.s. |
| Norvaline | 0.40 | 0.15 | 6.00 | 0.00 | 0.00 | 9.00 | -0.40 | 6.55 | 0.00 | ** |
| Ornithine | 0.00 | 0.00 | 6.00 | 0.21 | 0.35 | 9.00 | 0.21 | -1.84 | 0.10 | n.s. |
| Phenylalanine | 0.05 | 0.03 | 6.00 | 0.03 | 0.06 | 9.00 | -0.01 | 0.62 | 0.55 | n.s. |
| Proline | 0.00 | 0.00 | 6.00 | 0.49 | 0.49 | 9.00 | 0.49 | -2.97 | 0.02 | * |
| Serine | 0.93 | 0.53 | 6.00 | 0.08 | 0.19 | 9.00 | -0.85 | 3.79 | 0.01 | ** |
| Taurine | 0.00 | 0.00 | 6.00 | 0.15 | 0.18 | 9.00 | 0.15 | -2.51 | 0.04 | * |
| Threonine | 1.30 | 0.58 | 6.00 | 0.36 | 0.31 | 9.00 | -0.93 | 3.62 | 0.01 | ** |
| Tryptophan | 0.01 | 0.02 | 6.00 | 0.00 | 0.00 | 9.00 | -0.01 | 1.58 | 0.17 | n.s. |
| Tyrosine | 0.21 | 0.03 | 6.00 | 0.06 | 0.11 | 9.00 | -0.14 | 3.77 | 0.00 | ** |
| Valine | 4.65 | 3.92 | 6.00 | 45.90 | 2.53 | 9.00 | 41.25 | -22.80 | 0.00 | *** |

Supplementary table 2: **Numerical values of** **Complete amino acid composition before and after trace element optimization**

Supplementary figure 2: **Sum of coefficients of variation (CV) for each process parameter setting tested in iDOE 1**. Same colors indicate duplicates, showing reproducibility of setting 1 & 12, 2 & 13, 3 & 14 and 7 & 15. Setting 5 & 8 and 6 & 11 also resulted in low a sum of CVs, but only in one the duplicates and therefore they were excluded from the selection of stable process conditions. pH=7, T = 60°C resulted reproducibly in lowest sum of CVs.

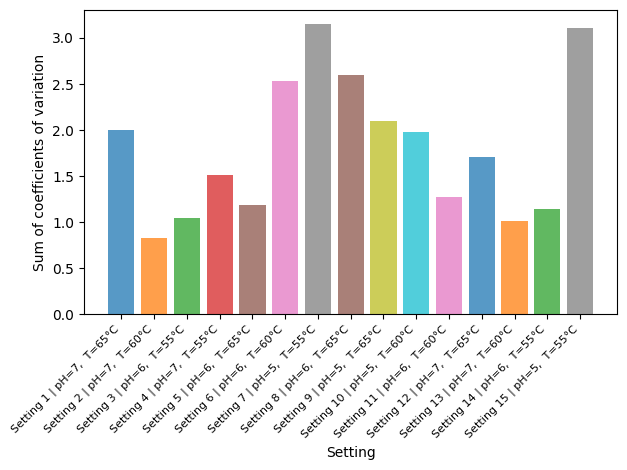

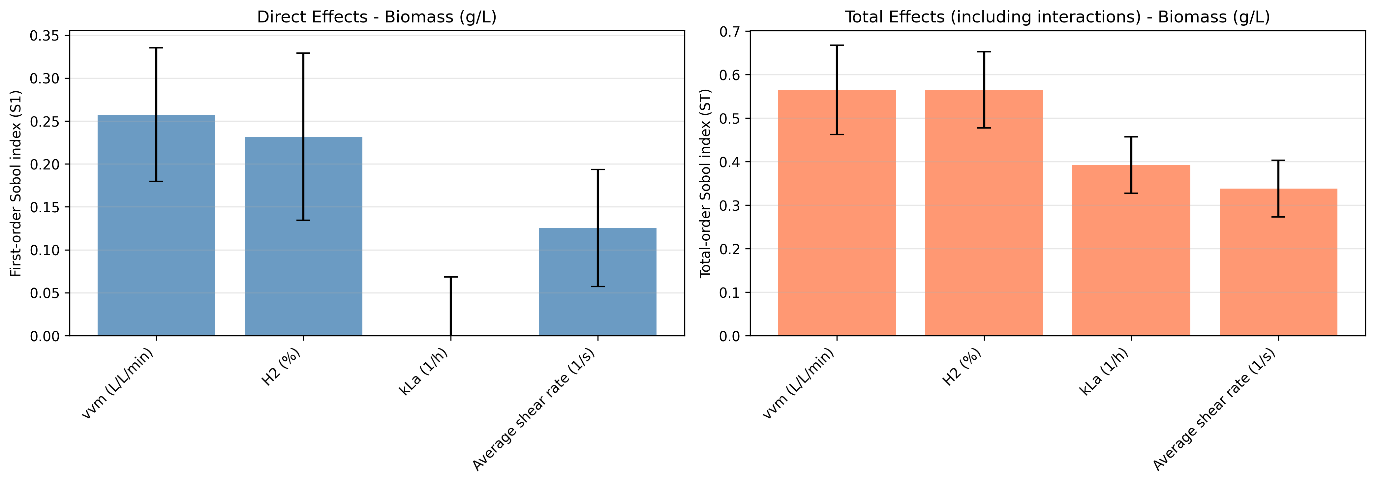

Supplementary figure 3: **First-order (S_1_) and total-order (S_t_) Sobol sensitivity indices for biomass concentration predicted by the iDOE hybrid model of iDOE 2 (n=512).** S_1_ indices quantify the direct contribution of each input variable to output variance, while Sₜ indices include both direct and interaction effects. vvm and H_2_ fraction show strong direct effects (S_1_ = 0.26, S_t_ = 0.57, S_1_ = 0.23,
S_t_ = 0.57), whereas k_L_a exhibits a negligible first-order effect but a substantial total-order contribution (S1 < 0.00, St = 0.39), indicating that its influence on biomass is primarily interaction-driven. Average shear rate also contributes to biomass formation (S_1_ = 0.13, S_t_ = 0.34), albeit to a lesser extent.

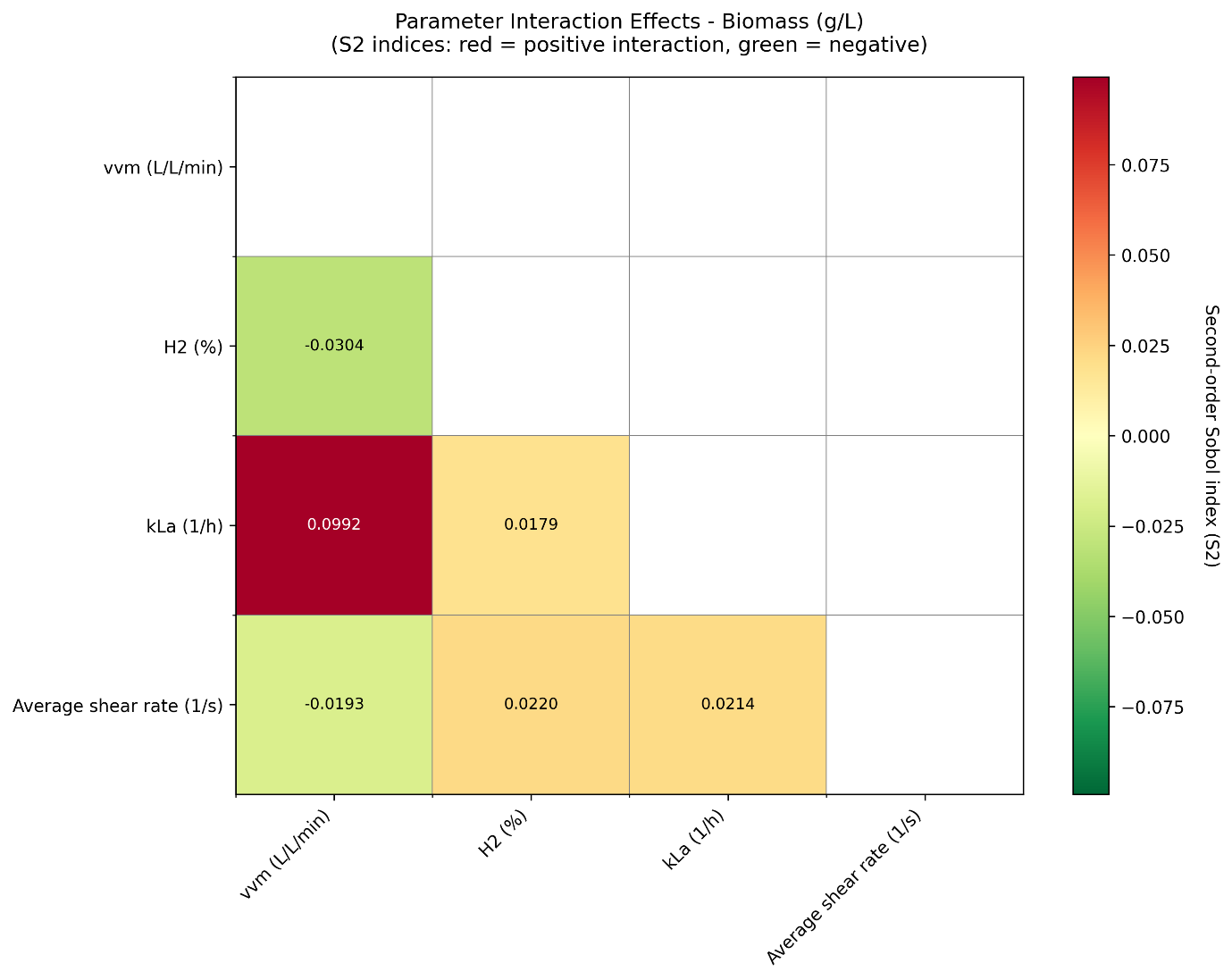

Supplementary figure 4: **Heatmap of second-order Sobol indices (S_2_) for biomass concentration predicted by the iDOE hybrid model of iDOE 2, showing pairwise interaction effects between input variables (n=512).** The strongest interaction is observed between vvm and k_L_a, consistent with the physical coupling of gas flow rate and volumetric mass transfer. Additional interactions involving shear rate and H_2_ fraction contribute to the overall variance but are of smaller magnitude.

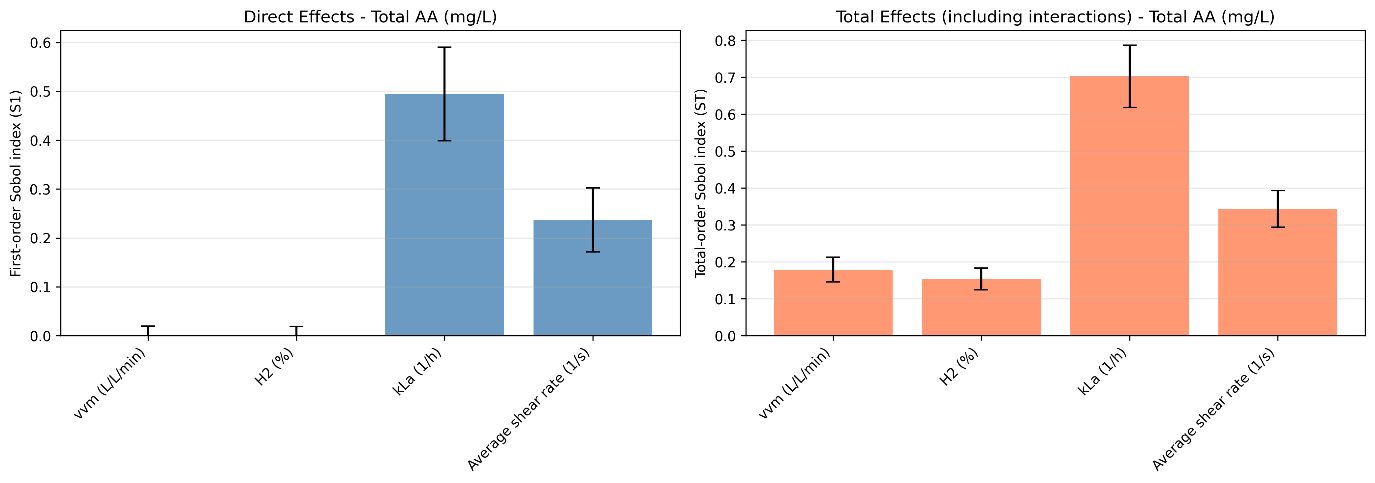

Supplementary figure 5: **First-order (S_1_) and total-order (S_t_) Sobol sensitivity indices for total amino acid concentration predicted by the iDOE hybrid model of iDOE 2 (n=512).** k_L_a dominates the response with the largest direct and total effects (S_1_ = 0.49, S_t_ = 0.70), followed by average shear rate (S_1_ = 0.24, S_t_ = 0.34). vvm and H_2_ fraction show minimal first-order effects (S_1_ < 0.00) but non-negligible total-order indices (S_t_ = 0.15 – 0.18),, indicating that their contribution to amino acid formation arises predominantly through interactions with other process variables.

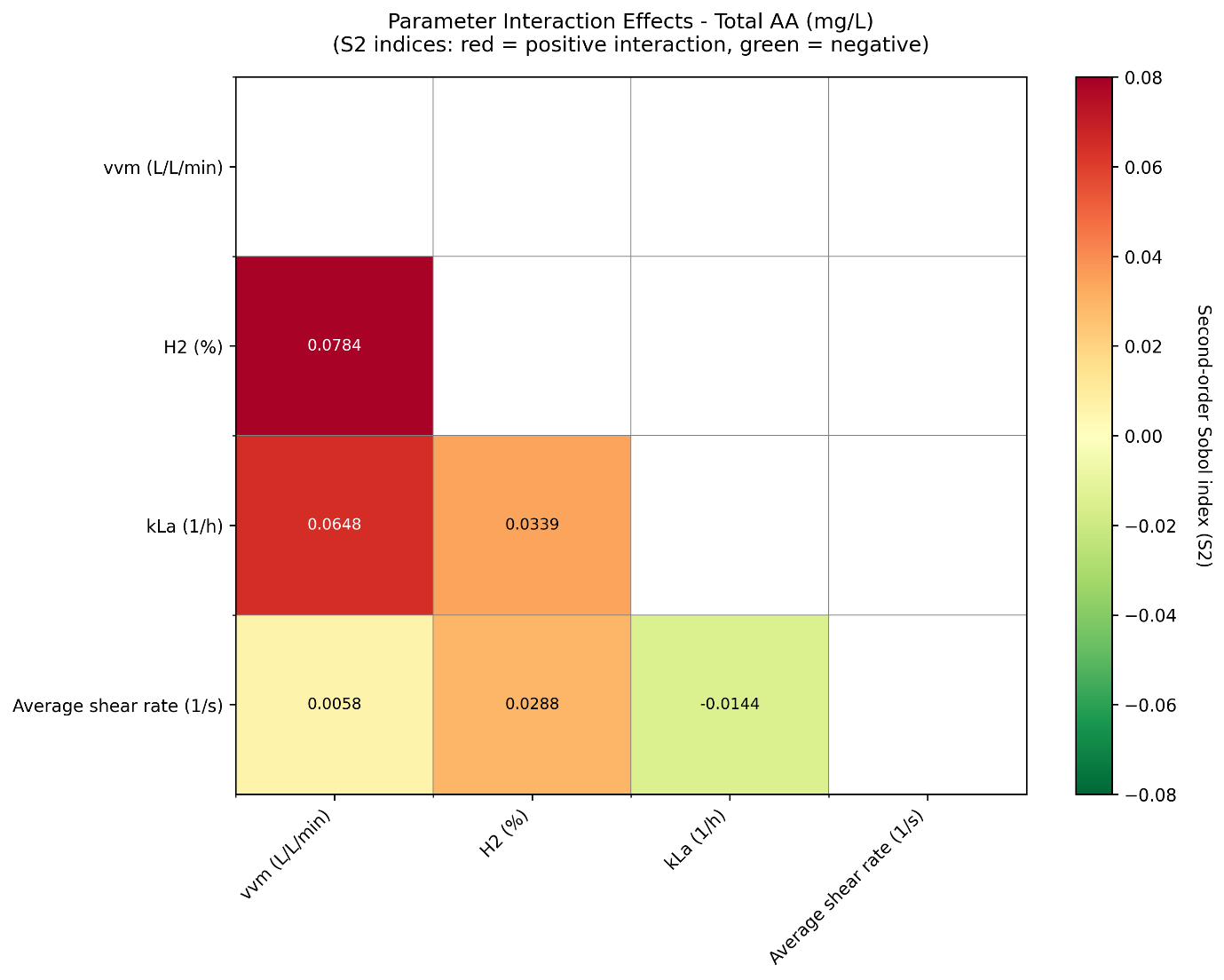

Supplementary figure 6: **Heatmap of second-order Sobol indices (S_2_) for total amino acid concentration predicted by the iDOE hybrid model of iDOE 2 (n=512).** Strong interaction effects are observed for combinations involving vvm, k_L_a, and H_2_ fraction, indicating that gas flow, gas composition, and mass transfer jointly govern amino acid production. These interaction-driven contributions explain the difference between first- and total-order effects observed in Supplementary figure 5.

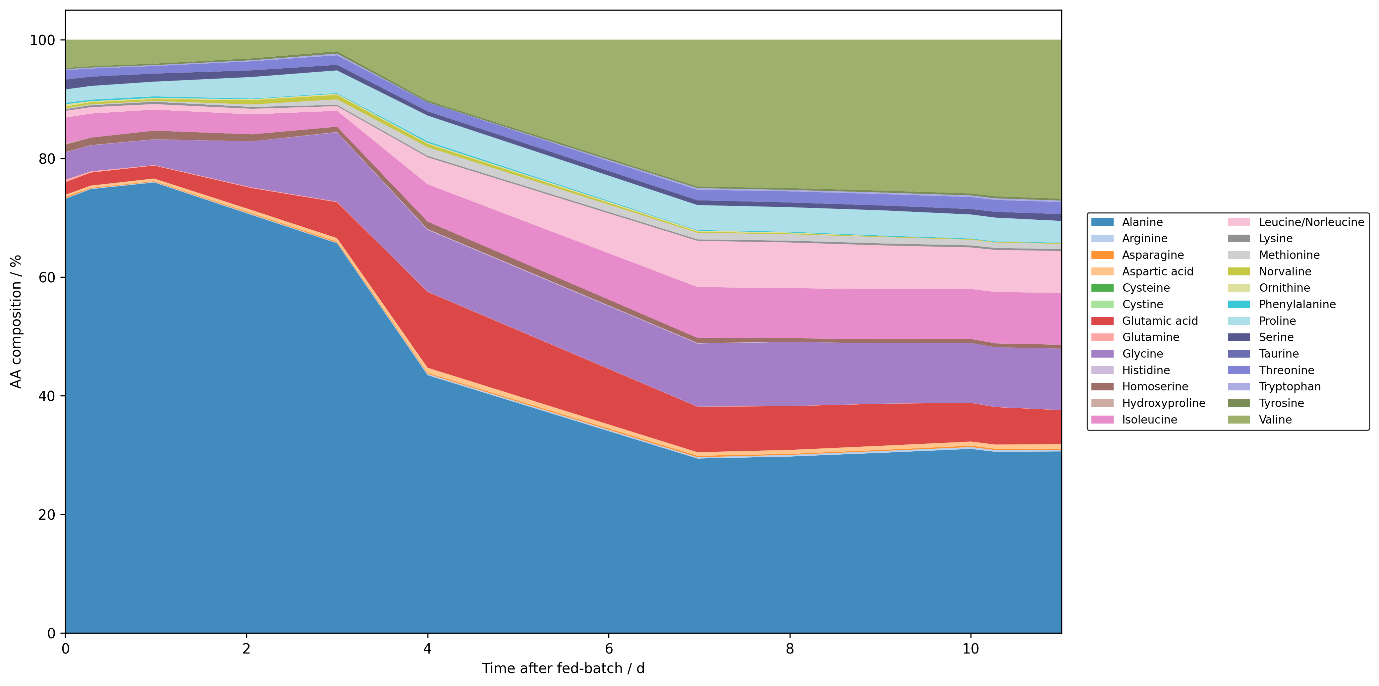

Supplementary figure 7: **Time-resolved amino acid composition during pilot-scale cultivation.** Relative amino acid (AA) composition during the 11-day pilot-scale (150 L) experiment. The stacked area plot illustrates the temporal evolution of the complete AA profile throughout the process.

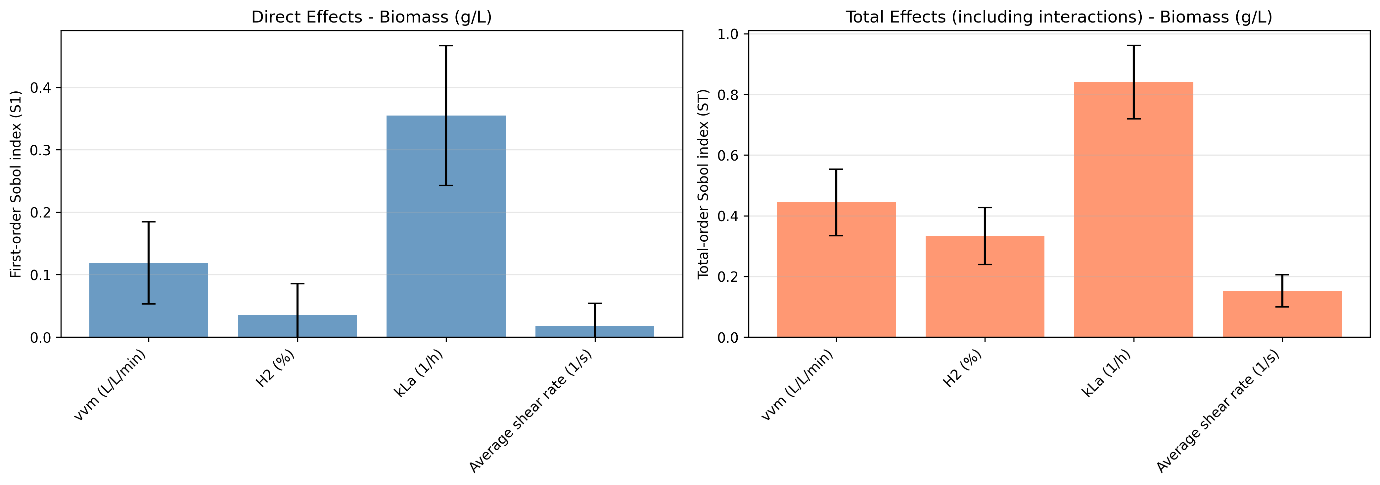

Supplementary figure 8: **First-order (S_1_) and total-order (S_t_) Sobol sensitivity indices for biomass concentration predicted by the steady-state hybrid model of iDOE 2 (n=512).** S_1_ indices quantify the direct contribution of each input variable to output variance, while Sₜ indices include both direct and interaction effects. In the steady-state model, k_L_a was identified as the dominant contributor to biomass formation (S_1_ = 0.35, S_t_ = 0.84), followed by vvm (S_1_ = 0.12, S_t_ = 0.44) and H_2_ fraction (S_1_ = 0.04, S_t_ = 0.33). Average shear rate exhibited only minor contribution (S_1_ = 0.02, S_t_ = 0.15). The larger total-order than first-order indices indicate that interaction effects contribute to biomass formation, according to the steady-state hybrid model.

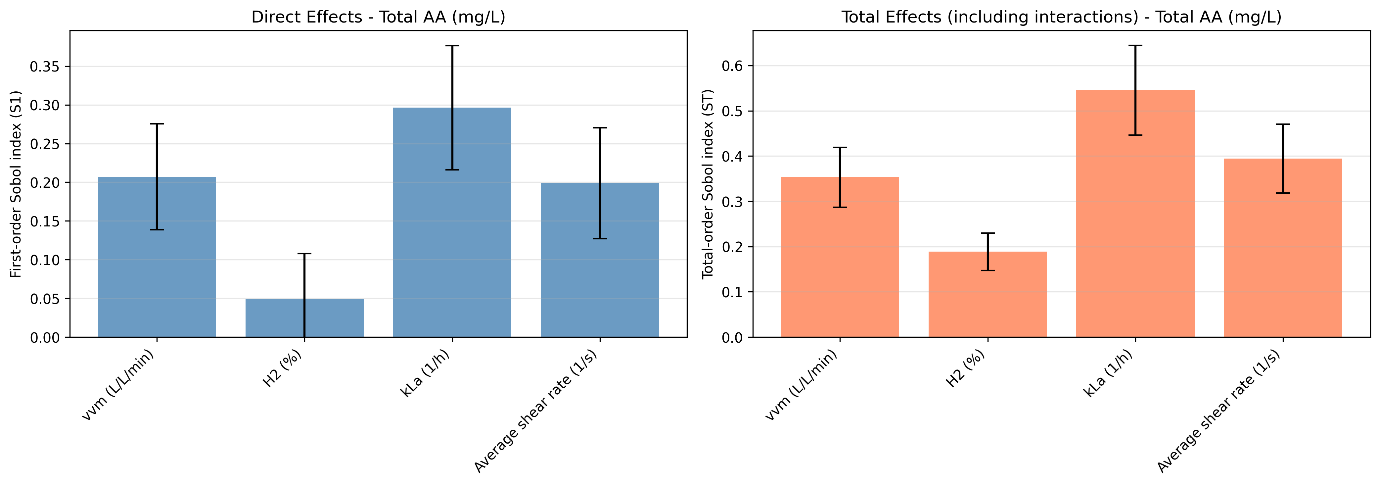

Supplementary figure 9: **First-order (S_1_) and total-order (S_t_) Sobol sensitivity indices for total amino acid concentration predicted by the steady-state hybrid model of iDOE 2 (n=512).**

The parameter k_L_a was identified as the dominant contributor to total AA production (S_1_ = 0.3, S_t_ = 0.55), followed by vvm (S_1_ = 0.21, S_t_ = 0.35), and average shear rate (S_1_ = 0.2, S_t_ = 0.39). H_2_ fraction exhibited only a minor contribution (S_1_ = 0.05, S_t_ = 0.19). The consistently larger total-order than first-order indices indicate that interaction effects contribute to total AA production.

Supplementary table 3: **Reduced media recipe for *M. marburgensis* cultivation**

| Compound | Concentration |
| --- | --- |
| NH_4_Cl (g L^−1^) | 2.10 |
| KH_2_PO_4_ (g L^−1^) | 6.80 |
| Trace Elements 1000x (mL L^−1^) | 1.00 |

Supplementary table 4: **DOE for trace element solution optimization**

| **Concentration in stock solution  (− … low, + … high, 0 … center)** | | | | |  |  |
| --- | --- | --- | --- | --- | --- | --- |
| **Cobalt** | **Iron** | **Nickel** | **Molybdenum** | **Magnesium** | **Repetitions** | **Experimental Design** |
| − | − | − | + | + | 3 | Fractional factorial design |
| + | − | − | − | − | 3 |  |
| − | + | − | − | + | 3 |  |
| + | + | − | + | − | 3 |  |
| − | − | + | + | − | 3 |  |
| + | − | + | − | + | 3 |  |
| − | + | + | − | − | 3 |  |
| + | + | + | + | + | 3 |  |
| 0 | 0 | 0 | 0 | 0 | 4 | Centerpoint |
| 0 | 0 | 0 | 0 | 0 | 3 | Negative control |
| − | − | − | − | − | 3 | Additional runs - part of full factorial design |
| − | + | − | − | − | 3 |  |
| + | + | − | − | − | 3 |  |
| − | − | + | − | − | 3 |  |
| + | − | + | − | − | 3 |  |
| + | + | + | − | − | 3 |  |
| − | − | − | + | − | 3 |  |
| + | − | − | + | − | 3 |  |
| − | + | − | + | − | 3 |  |
| + | − | + | + | − | 2 |  |
|  |  |  |  |  | 60 | Σ Runs |

*Fractional factorial screening design with 5 factors and 2 levels, including centerpoint runs, negative controls without inoculation and additional runs part of the full factorial design for ideal utilization of all available serum bottles. The level “+” indicates a higher, “**−** “ a lower and “0” the same concentration as in the original media recipe. The exact trace element composition is part of an undisclosed process and therefore cannot be reported in absolute terms.

pH / -

| Temperature / °C | R1 |
| --- | --- |
| 65 | 7 |
| 60 | 7 |
| 55 | 6 |
| 55 | 7 |

| Temperature / °C | R2 |
| --- | --- |
| 65 | 6 |
| 60 | 6 |
| 55 | 5 |

| Temperature / °C | R3 |
| --- | --- |
| 65 | 6 |
| 65 | 5 |
| 60 | 5 |
| 60 | 6 |

| Temperature / °C | R4 |
| --- | --- |
| 65 | 6.95 |
| 60 | 6.95 |
| 55 | 5.95 |
| 55 | 5 |

Supplementary figure 10: **Setup of iDOE1**: Two-factor, 3-level full factorial intensified design across 4 bioreactors used in iDOE1 for finding most stable conditions regarding temperature and pH. The numbers indicate the unique setpoint number of the process parameter combination.

|  | | | | 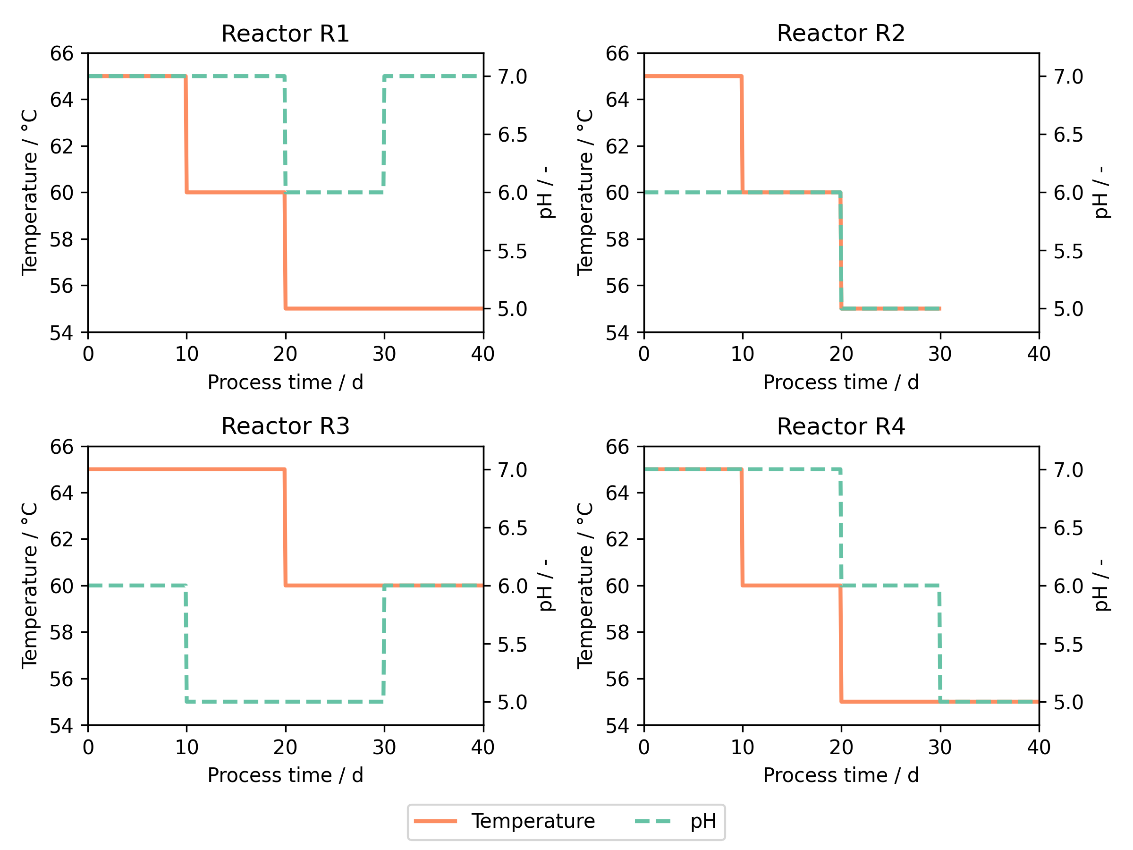 |
| --- | --- | --- | --- | --- |
| **Setting** | **T / °C** | **pH** | **R** |  |
| 1 | 65 | 7 | R1 |  |
| 2 | 60 | 7 |  |  |
| 3 | 55 | 6 |  |  |
| 4 | 55 | 7 |  |  |
| 5 | 65 | 6 | R2 |  |
| 6 | 60 | 6 |  |  |
| 7 | 55 | 5 |  |  |
| 8 | 65 | 6 | R3 |  |
| 9 | 65 | 5 |  |  |
| 10 | 60 | 5 |  |  |
| 11 | 60 | 6 |  |  |
| 12 | 65 | 7 | R4 |  |
| 13 | 60 | 7 |  |  |
| 14 | 55 | 6 |  |  |
| 15 | 55 | 5 |  |  |

Supplementary figure 11: **Time-resolved process parameters for iDOE 1**. Each reactor (R1–R4) was operated with dynamic shifts in temperature and pH setpoints over the course of a single experiment. The left table lists all applied setpoints and their allocation. The corresponding profiles on the right show the stepwise changes in temperature (orange solid line) and pH (green dashed line) over the process time in d, for each reactor.

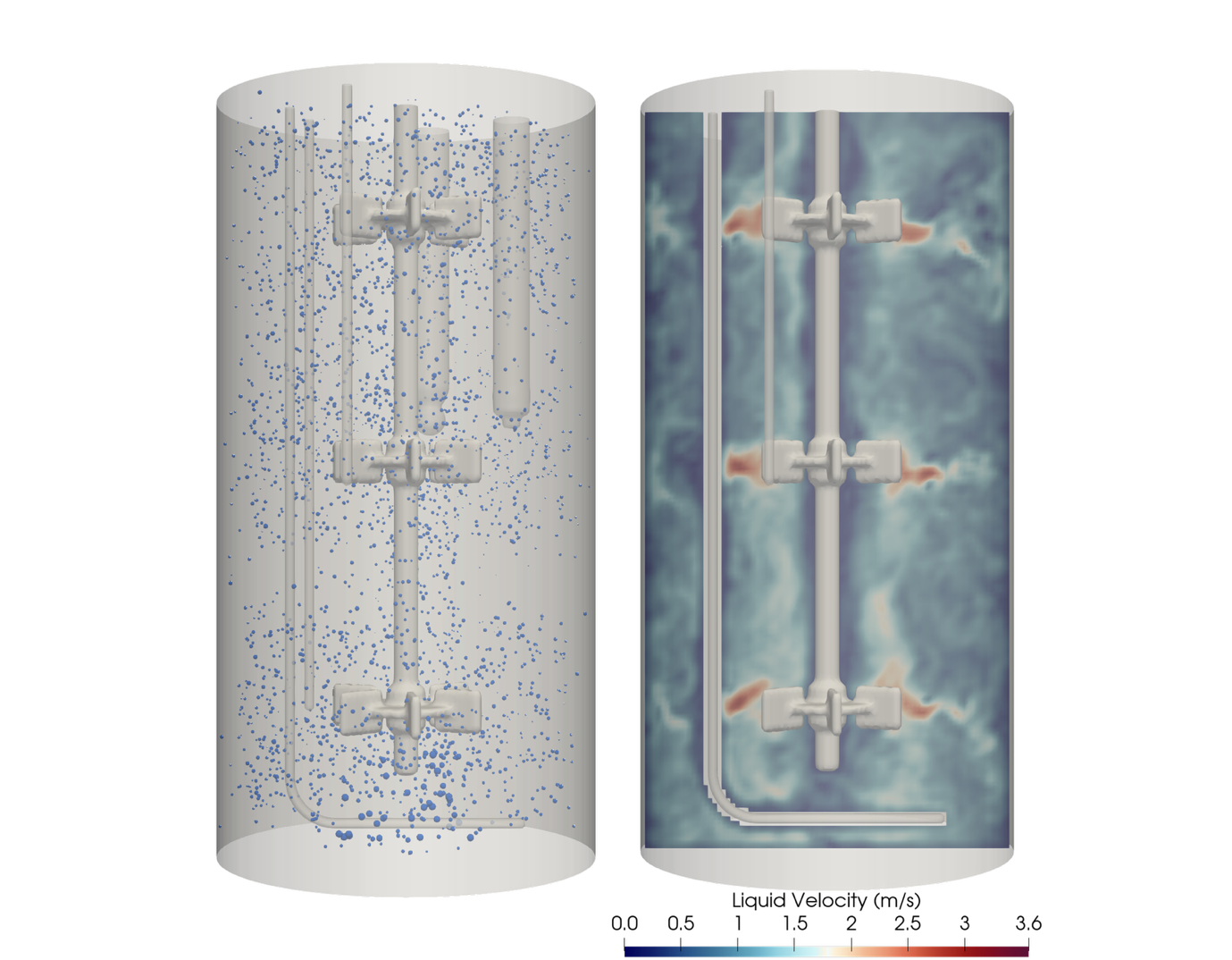

Supplementary figure 12: **Visualization of bubbles and velocity field from CFD results of the Eppendorf DASGIP at 1200 rpm and 0.1875 vvm.**

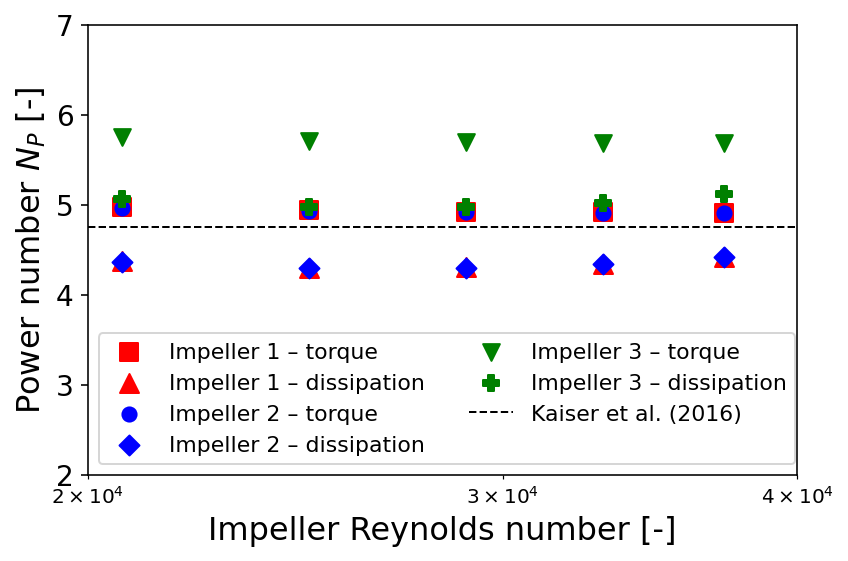

Supplementary figure 13: **Impeller power number per stage over Reynolds number and comparison to literature value from Kaiser *et al*.**^1^**.**

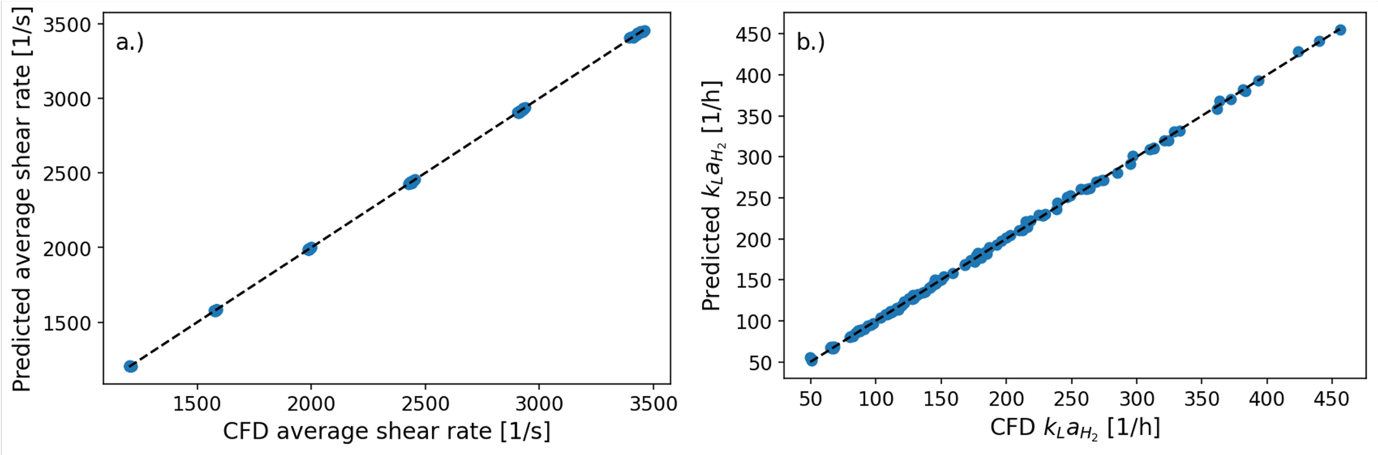

Supplementary figure 14: **Figure 3: Parity plots of machine learning model for average shear rate (a) and k_L_aH_2_ (b).**

1.	Kaiser, S. C., Werner, S., Jossen, V., Kraume, M. & Eibl, D. Development of a method for reliable power input measurements in conventional and single-use stirred bioreactors at laboratory scale. *Eng. Life Sci.* **17**, 500–511 (2017).
